# A ratchet-like apical constriction drives cell ingression during the mouse gastrulation EMT

**DOI:** 10.1101/2022.04.30.489707

**Authors:** Alexandre Francou, Kathryn V. Anderson, Anna-Katerina Hadjantonakis

**Affiliations:** Developmental Biology Program, Sloan Kettering Institute, Memorial Sloan Kettering Cancer Center, New York, NY 10065, USA

## Abstract

Epithelial-to-Mesenchymal Transition (EMT) is a fundamental process in which epithelial cells acquire mesenchymal phenotypes and the ability to migrate. EMT is the hallmark of gastrulation, an evolutionarily conserved developmental process. In mammals, epiblast cells lose pluripotency and ingress at the primitive streak to form mesoderm. Cell exit from the epiblast epithelial layer and the associated EMT are dynamically regulated processes involving a stereotypical sequence of cell behaviors. 3D time-lapse imaging of gastrulating mouse embryos combined with cell and tissue scale data analyses revealed the stochastic-like ingression of epiblast cells at the primitive streak. Ingressing cells constrict their apical surfaces in a pulsed ratchet-like fashion through asynchronous shrinkage of apical junctions. A quantitative analyses of the distribution of apical proteins, revealed the anisotropic and complementary distribution of members of the actomyosin network and Crumbs2 complexes, potential regulators of asynchronous shrinkage of cell junctions. The analysis of mutants demonstrated a requirement for Crumbs2 in Myosin2 localization and activity at apical junctions, and as a candidate for regulating actomyosin anisotropy.

## INTRODUCTION

Epithelial-to-Mesenchymal Transitions (EMTs) are morphogenetic programs necessary for the formation of tissues throughout embryonic development, cancer progression and metastasis (Francou and Anderson, 2020; Thiery, 2002; Thiery et al., 2009; Ye and Weinberg, 2015). The EMT that occurs during embryo gastrulation is an evolutionarily conserved event occurring in triploblasts as they form three definitive germ layers (Lim and Thiery, 2012; Nakaya and Sheng, 2008; Thiery et al., 2009).

In amniotes the onset of gastrulation involves formation of the primitive streak at the posterior epiblast, at around embryonic **(E)** 6.5 day in the mouse (Ferrer-Vaquer et al., 2010; Nakaya and Sheng, 2008; Williams et al., 2012) (Figure 1A). Primitive streak initiation is regulated by the convergence of WNT, BMP and Nodal signals, which together with an FGF input trigger the EMT process (Ferrer-Vaquer et al., 2010; Morgani and Hadjantonakis, 2020; Ramkumar and Anderson, 2011). The basement membrane is broken down at the primitive streak facilitating movement of cells out of the epiblast tissue layer, accompanied by changes in their cell shape, including apical constriction and basal translocation of cell bodies (Ramkumar et al., 2016; Williams et al., 2012). Basal cell ingression and migration away from the epiblast epithelium is facilitated though the dissolution of apical junctions. As cells at the primitive streak undergo EMT, more laterally positioned epiblast cells converge toward the midline (Williams et al., 2012), replenishing the pool for continued ingression and ensuring epithelial integrity.

**Figure 1:**
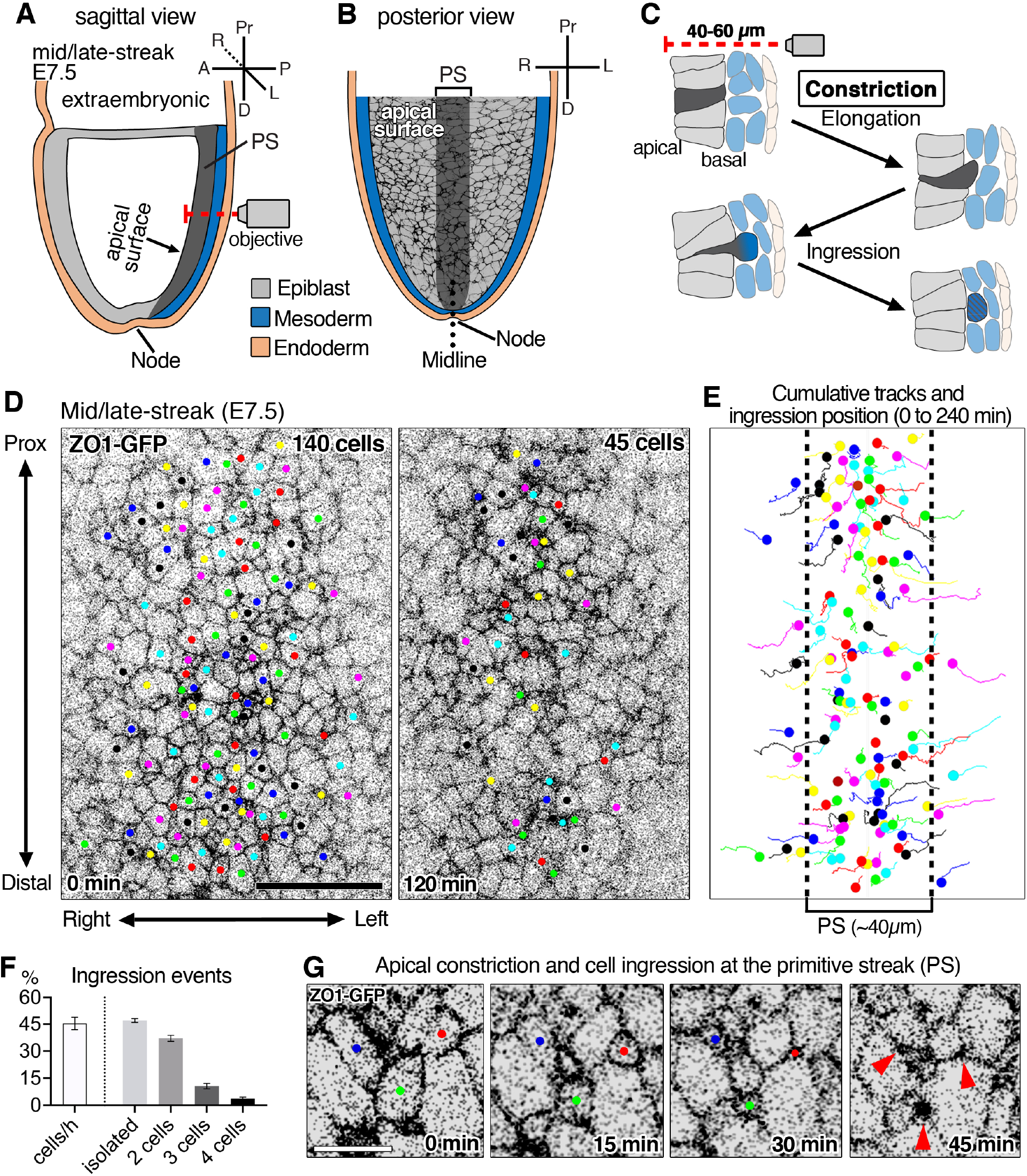
Time-lapse imaging of ZO-1-GFP reporter reveals EMT events at the primitive streak of the mouse gastrula and apical constriction associated with cell ingression. **(A)** Schematic sagittal section view of E7.5 mouse embryo and 3D time-lapse imaging performed from the posterior side with an objective positioned adjacent to the primitive streak (PS). Several tissue layers need to be imaged across (including visceral endoderm and mesoderm) to reach the epiblast apical surface situated furthest from the objective (red dashed line shows the microscope light path). **(B)** Schematic of a view of the epiblast layer apical surface from the inner cavity. The midline (dotted line axis) separates the right and left sides of the embryo. **(C)** High magnification schematic of a sagittal view of an EMT event at the primitive streak depicting a cell constricting its apical surface, elongating basally (dark grey), and ingressing out of the epiblast layer to integrate the mesoderm (blue). Note the apical surface of epiblast cells is located 40-60μm away from the microscope objective. **(D)** Single time-points at t=0min and t=120min of a time-lapse of a ZO-1-GFP embryo. Movies were analyzed as maximum projections of z-stacks to visualize the apical surfaces of cells. Here all the cells that can be followed and observed ingressing during 4hr were tracked. Of 140 cells initially tracked, 95 cells constricted and ingressed over the course of 120min (45 tracked cells remaining). (Time lapse with 5min time interval, 290 cells in the field of view, 140 ingressing cells tracked). **(E)** Cumulative tracking of the 140 cells showing cell tracks over time (lines) and cell position during ingression event (dots). Epiblast cells converge toward the primitive streak (~40μm, dotted lines) where the majority of ingression events occur. **(F)** Graphs showing 44±2.1% of cells in the primitive streak ingress each hour, with 48±1.1% ingressing as single cells, 37±1.7% as pairs of cells, 11±1.5% as triplets (groups of 3 cells), and 4% as groups of 4 cells (of a total of 378 ingressing cells analyzed in 3 embryos). **(G)** High magnification view showing the apical constriction of 3 cells at the primitive streak. Pr: proximal, D: distal, A: anterior, P: posterior, R: right, L: left, Ext Emb: extra embryonic region, PS: primitive streak. Error bars represent s.e.m. Scale bars, d, 40μm; g, 10μm. **Figure supplement 1**: Apical constriction and ingression of epiblast cells in the vicinity of the mouse primitive streak.

Apical constriction is an epithelial cell shape change associated with morphogenetic processes such as tissue bending, cell delamination and internalization (An et al., 2017; Chung et al., 2017; Lecuit and Lenne, 2007; Martin and Goldstein, 2014; Nishimura et al., 2012; Samarage et al., 2015; Simoes et al., 2017). The dynamics of apical constriction at gastrulation have been studied in invertebrates, particularly in Drosophila, and shown to be controlled by apical actomyosin contractility (Marston et al., 2016; Martin and Goldstein, 2014; Martin et al., 2009; Mason et al., 2013; Roh-Johnson et al., 2012). Apical constriction has been associated with cell ingression during the gastrulation EMT in chick and mouse (Rozbicki et al., 2015; Serrano Najera and Weijer, 2020; Williams et al., 2012), but how cells dynamically constrict apical surfaces during EMT, and what regulates this process remain open questions. Gastrulation process in Drosophila and mouse cannot be directly compared as they exhibit major different time-scales (20min vs. >24hr days), mechanisms and spatial parameters (tissue invagination vs. cell ingression). In the chick embryo, Myosin-2 plays a role in the formation of the primitive streak and cell ingression from the epiblast layer (Chuai et al., 2006; Rozbicki et al., 2015; Serrano Najera and Weijer, 2020; Voiculescu et al., 2007; Voiculescu et al., 2014). Crumbs2, a protein shown to regulate apical polarity in epithelia, is critical for cell ingression during gastrulation EMT in mouse embryos (Ramkumar et al., 2016). Apical Crumbs2 and Myosin-2B show an anisotropic accumulation in mouse epiblast cells in the vicinity of the primitive streak (Ramkumar et al., 2016), reminiscent of patterns controlling apical constriction during Drosophila salivary gland development (Roper, 2012), suggesting these proteins are potential regulators of apical constriction during mouse EMT. Visualizing the cellular dynamics of the epiblast during mouse gastrulation has been a longstanding challenge due to its internal location which limits optical access and visualization at high resolution.

Using live imaging of *ex utero* cultured mouse embryos, and high resolution visualization and segmentation of epiblast cell membranes, we performed a dynamic quantitative analysis of apical constriction associated with ingression during the mouse gastrulation EMT. We observed cells undergoing EMT at the mouse primitive streak in a stochastic-like manner, constricting their apical surfaces in a rachet-like pulsed fashion through asynchronous shrinkage of apical cell-cell junctions. By analyzing and quantifying the distribution of apical proteins, we uncovered the anisotropic and complementary distribution of actomyosin network and Crumbs complex proteins consistent with a role in regulating junctional shrinkage and apical constriction. The localization of the apical actomyosin network, as well as two kinases aPKC and Rock1, was perturbed in *Crumbs2* mutants, thereby identifying key components of a putative regulatory network driving apical constriction.

## RESULTS

### Epiblast cells undergo apical constriction and isolated ingression at the mouse primitive streak

The mouse gastrulation EMT occurs at the primitive streak, which, by contrast to the chick, does not form a morphologically distinct domain (Figure 1A,B). Columnar epithelial epiblast cells with their apical surfaces facing the inner (amniotic) cavity of the embryo apically constrict and elongate basally as they ingress out of the epiblast layer (Figure 1A,C) (Williams et al., 2012). To visualize the dynamic changes in the shapes of all cells in tissue context at single cell during the gastrulation EMT, we sought to label the membranes of all epiblast cells and perform time-lapse imaging. Time-lapse imaging of the epiblast cells in the vicinity of the primitive streak of developing mouse embryos is challenging for several reasons. Embryos need to be kept intact as tissues cannot be microdissected, the epiblast tissue is cup-shaped with inherent curvature and located within the deepest extremity of the embryo (up to 60μm away from the objective), needing to be imaged through the visceral endoderm and mesoderm tissue layers (Figure 1C). Even light-sheet systems, which by merging multi-view data can provide *in toto* visualization of cells in developing embryos, have predominantly been used with nuclear-localized reporters or scattered membrane-labelled cells, data which are less challenging to segment than widespread membrane-localized reporters (McDole et al., 2018).

We used a ZO-1-GFP protein fusion reporter (Foote et al., 2013) to visualize the tight junction and the apical surface of epiblast cells in order to quantify apical surface dynamics (Figure 1D); and a membrane-localized *R26^mT/mG^* reporter to visualize their entire plasma membrane and identify completion of apical constriction associated with cell ingression out of the epithelial layer (Figure 1-figure supplement 1A and Video S1). 3D time-lapse imaging of embryos at mid/late-streak stage (E7.5) was performed to image the primitive streak) (Figure 1A, B). Time-lapse imaging of ZO-1-GFP embryos revealed that epiblast cells undergo extensive rearrangements as cells on the left and right sides of the embryo converge towards the primitive streak (Figure 1D, Figure 1-figure supplement 1C, Video S2,3) (Williams et al., 2012). The majority of epiblast cells underwent apical constriction and ingression within a ~40μm region in the posterior midline (Figure 1D,E, Video S2), corresponding to the domain of Snail expression and basement membrane breakdown within the epiblast (Francou and Anderson, 2020; Ramkumar et al., 2016; Williams et al., 2012), which by convention defines the primitive streak (Figure 1E dotted lines). Rare apical constriction and ingression events were also observed outside the domain of the primitive streak (Figure 1-figure supplement 1B and Video S3 bottom).

Within a one hour period 44±2% of cells within the primitive streak domain, constricted and ingressed (mean±s.e.m., n=378 cells, 3 embryos) (Figure 1D,E,F, Video S2), with ingression events appearing to occur stochastically (3-4h). Among the ingressing cells tracked, 48% constricted and ingressed as isolated events (more than 30min apart from adjacent neighboring cells), while the remaining 52% ingressed as pairs, and occasionally as groups of 3-4 cells (less than 30min apart from adjacent neighboring cells) (Figure 1F). At the tissue level the epiblast underwent extensive rearrangement, with some cells moving and acquiring new neighbors prior to their coordinate ingression (Figure 1-figure supplement 1D,E), while others, which were initially neighbors, ingressed at different times (more than 30 min apart) (Figure 1-figure supplement 1D,F).

### Epiblast cells undergo a ratchet-like pulsed apical constriction during ingression

We next focused on changes associated with the apical surfaces of cells during their ingression. Apical constriction has been extensively studied in Drosophila, including during mesoderm formation and neuroblast delamination (An et al., 2017; Martin et al., 2009; Mason et al., 2013; Simoes et al., 2017), but few studies have investigated the gastrulation EMT in an amniote, and it is unknown how cells constrict their apical surfaces during single cell ingression in the mouse embryo versus the more global constriction and invagination happening during Drosophila gastrulation. To follow changes in apical surfaces during cell ingression, cell membranes were segmented using Tissue Analyzer software (Aigouy et al., 2010; Aigouy et al., 2016) on projections of z-stacks from time-lapse data. By visualizing cells at the primitive streak, we identified apical constriction associated with cell ingression and quantified several parameters, including apical surface area, apical cell surface elongation and orientation, and number of apical junctions (Figure 1G, Figure 2,3-figure supplement 1, Video S4,5). Epiblast cells exhibited a variety of apical surface sizes and shapes that fluctuated over time as they flowed toward the primitive streak. As they initiated apical constriction at the primitive streak, cells exhibited 20-80μm^2^ apical surface area, and completed ingression within 25-90min (Figure 2A and Figure 2,3-figure supplement 1B). With the time-scale we are to image and keep living embryos and fluorescent signal integrity, majority of ingressing cells exhibited pulsed constrictions, generally consisting of 2-4 major phases of contraction (Figure 2B,C and Figure 2,3-figure supplement 1F). As cells progressed towards ingression, their number of apical junctions decreased (Figure 2,3-figure supplement 1J, Video S5).

**Figure 2:**
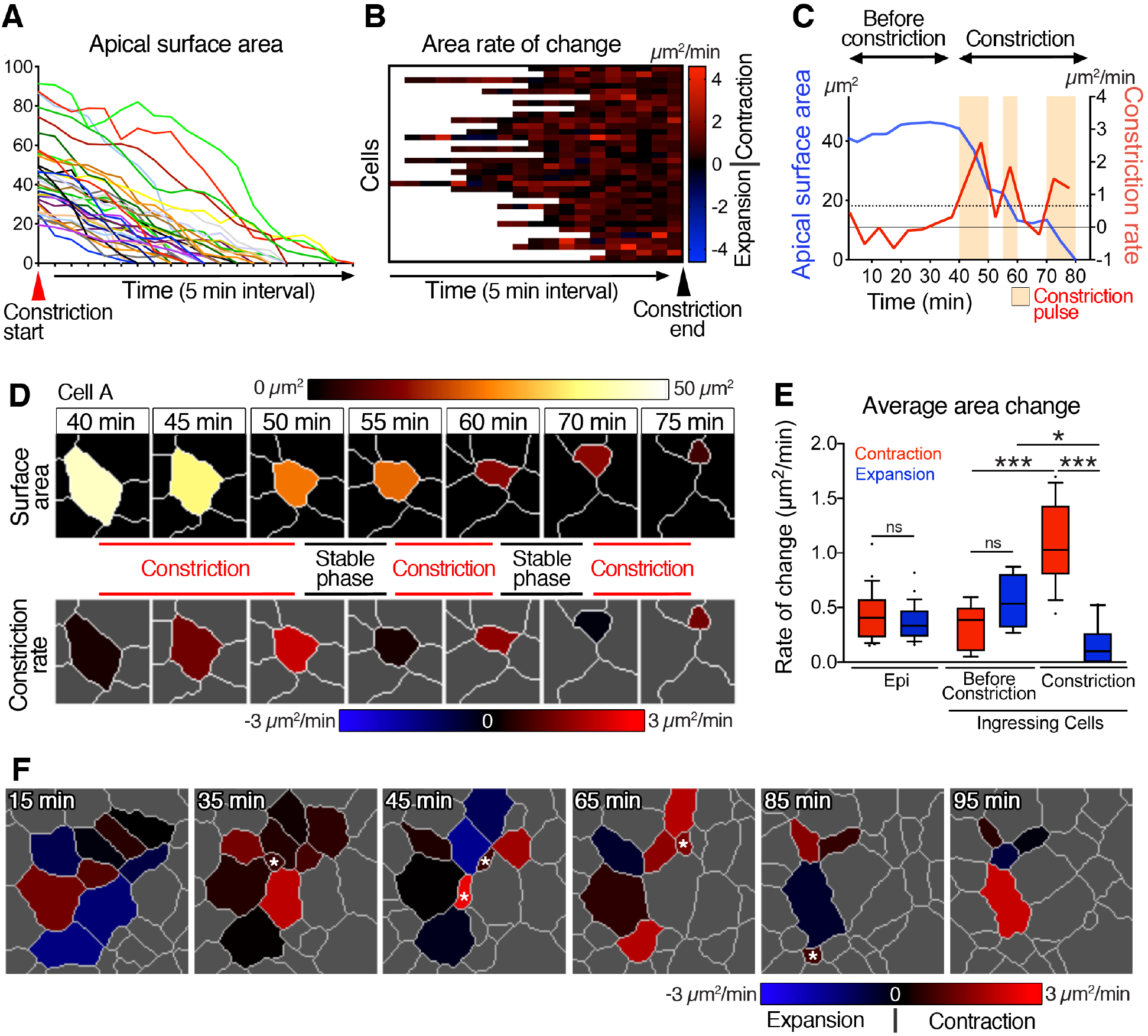
Pulsed racket-like apical constriction during epiblast cell ingression and asynchronous constriction at multi-cellular level. **(A)** Graph showing the apical surface area of a few cells during constriction associated with ingression, aligned to the start of constriction. **(B)** Heat map plot showing the rate of changes of apical surface area of cells during the final constriction period, revealing minimal expansion and pulsed contractions. Each row represents data for an individual cell and are aligned to the end of constriction. **(C)** Graph of apical surface area and rate of constriction of a single cell (cell A) showing slight oscillation in apical surface before undergoing a final constriction period with pulses of constriction, here with three main pulses (beige regions). **(D)** Membrane segmentation and color-coded time-series of apical surface area and constriction rate of a single cell (cell A) showing three pulses of constriction separated by stable steps. (cell corresponding to the plot showed in C). **(E)** Plot showing the average rate of area changes. Epiblast cells away from the primitive streak before ingression show low and equivalent levels of contraction and expansion. Cells at the primitive streak show no significant differences between contraction and expansion before the constriction, whereas rate of contraction is significantly higher, and expansion reduces during the subsequent constriction period. **(F)** Color-coded time-series and heat map plot of rate of change of apical surface area showing asynchronous oscillation in area and asynchronous constriction and ingression among neighbors. Asterisks show ingression events. 41 cells quantified from 4 embryos. Error bars represent s.e.m. *P<0.05, ***P<0.001 (Unpaired bilateral Mann-Whitney test). **Figure supplement 1**: Details and features of constriction pulses and cell parameters during constriction. **Figure supplement 2**: Asynchronous apical constriction at the multi-cellular level.

A closer examination of individual cells allowed us to quantify features of apical surface area throughout the constriction period. The example in Figure 2D depicts the apical surface area of a cell exhibiting 3 main pulses of constriction each separated by more stable intermediate steps (Figure 2C,D, Video S6). While cells showed an average rate of constriction of 1.1±0.1 μm^2^/min (mean±s.e.m., n=51 cells, 4 embryos), they exhibited constriction pulses with higher rates of contraction (up to 4.6 μm^2^/min), and an average pulse magnitude of 1.4±0.1 μm^2^/min (Figure 2B,C,E and Figure 2,3-figure supplement 1G). Cells at the primitive streak fluctuated between expansion and contraction before ingression, as did epiblast cells located some distance from the streak (Figure 2C and Figure 2,3-figure supplement 1B,E). During ingression, cells spent 90% of the time contracting apical surfaces (compare with 48% and 55% for epiblast cells away from the streak and before ingression) with an average rate of contraction of 1.1μm^2^/min, higher than the average expansion rate of 0.1μm^2^/min (Figure 2E and Figure 2,3-figure supplement 1D). To conclude, epiblast cells exhibited fluctuations in apical surface areas before ingression, and pulsed constrictions during ingression at the primitive streak. No hallmark or significant trend was observed at the onset of constriction, the elongation and orientation of the apical surface at the beginning or throughout the constriction period, nor the number of edges when constriction initiated (Figure 2,3-figure supplement 1J-L).

### Asynchronous apical constriction at multi-cellular level

We analyzed clusters of cells at the primitive streak to uncover details of the apical constriction behavior at the multi-cellular level. Segmenting cell clusters allowed tracking of cells and documentation of changes in their apical surfaces through time, as cells ingressed (Figure 2F, Figure 2-figure supplement 2, Video S7). Cells underwent fluctuations in apical surface area as some cells constricted and ingressed (Figure 2-figure supplement 2B, Video S7). By color-coding and plotting the rate of apical change, we observed asynchronous oscillation of expansion and contraction phases among neighboring cells. As cells constricted and ingressed, the neighbors did not always exhibit the same behavior; some contracted at the same time, while others slightly expanded or maintained a constant surface area (Figure 2F, asterisk show ingression; S3C and Video S7). Thus, neighboring epiblast cells undergoing EMT and ingressing at the primitive streak displayed asynchronous apical behaviors and ingress in a stochastic-like manner, with some neighboring cells ingressing 2 hours apart, in comparison to Drosophila gastrulation where all ventral furrow cells constrict in less than 10min.

### Asynchronous shrinkage of junctions correlates with apical constriction during cell ingression

Most apical junctions of any given cell decreased in length in a pulsed manner, with 1-3 pulses of shrinkage (Figure 3A,B), with junctions appearing to shrink asynchronously (Figure 3A,D, Video S6). By analyzing and quantifying the behavior of each junction of any one cell during constriction, we noted that the average rate of shrinkage was significantly higher during cell constriction pulses (Figure 3C). Junctions of individual cells shrunk asynchronously (Figure 3D and Figure 2,3-figure supplement 1H,I), differentially and with different magnitudes between one constriction pulse and the next (Figure 3E and Figure 2,3-figure supplement 1H,I), suggesting that asynchronous differential junctional shrinkage may trigger constriction of the cell surface. In sum, epiblast cells at the mouse primitive streak constricted their apical surfaces in a ratchet-like manner, with asynchronous shrinkage of junctions (Figure 3F).

**Figure 3:**
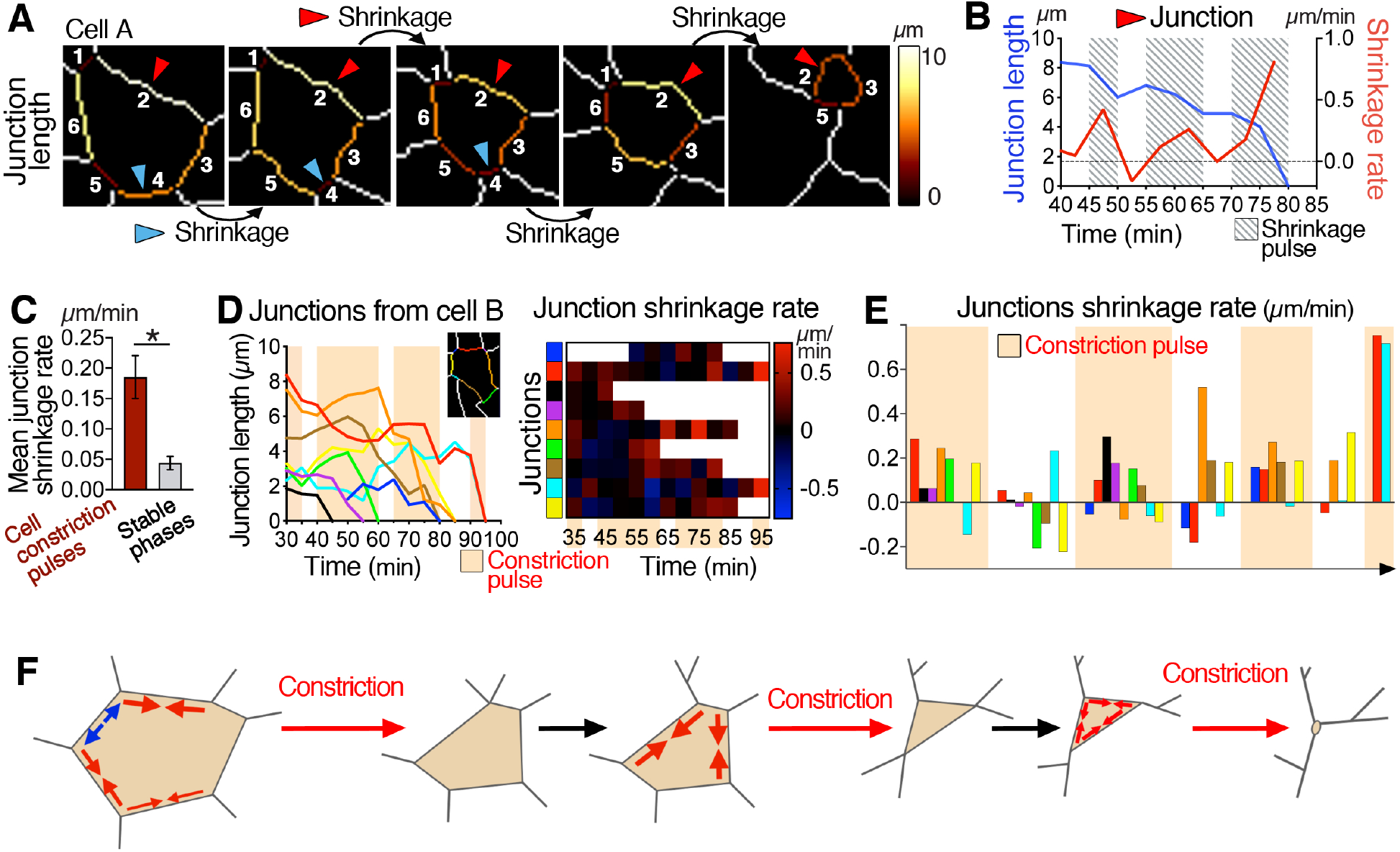
Asynchronous junctional shrinkage during apical constriction. **(A)** Membrane segmentation and color-coded time-series of single junction showing reduction of junctions length during apical constriction of cell A showed in Figure 2. Here the six initial junctions are numbered, and two are pointed (arrowheads) and shrink at different time. **(B)** Graph showing the length and rate of shrinkage of the junction identified with the red arrowhead in A, indicating the 3 main pulses of shrinkage (hatched regions). **(C)** Graph showing increased rate of junctional shrinkage during cell constriction pulses compare to stable phases between constriction pulses from multiple cells (3 embryos, 51 junctions). **(D)** Graph showing the length of individual junctions (left) during apical constriction of a single cell (cell B plotted in Figure 2,3-figure supplement 1A). Junctions reduce their length differentially during the constriction pulses (beige regions). Heat map plot of the shrinkage rate of junctions (right), showing junctions shrinking asynchronously during the constriction pulses (beige time points). Each row represents data for an individual junction, junctions are color-coded as corresponding in the left plot and ordered as neighboring junctions. **(E)** Plot of the average shrinkage rate of each junction through time during each constriction pulse and stables phases, showing the asynchronous shrinkage of different sets of junctions with variable magnitude during the different cell constriction pulses. Junctions are color-coded as neighbors as in D. **(F)** Model of rachet-like pulsed constriction associated with asynchronous junctional shrinkage. Error bars represent s.e.m. *P<0.05 (Unpaired bilateral Mann-Whitney test).

### Anisotropic accumulation of components at apical junctions of epiblast cells

We sought to analyze the localization of apical proteins implicated in epithelial apical constriction in different tissues and species, that could potentially be involved in the apical constriction associated with the mouse gastrulation EMT. E7.5 embryos were immunostained and epiblast apical surface imaged in regions proximo-distally middle of the primitive streak (least curved and best for analysis) (Figure 4-figure supplement 1A). Crumbs2 and Myosin2B have been described as exhibiting heterogenous and complementary distributions at the apical junctions of epiblast cells (Ramkumar et al., 2016), and our data showed Myosin2B and Crumbs2 were predominantly anisotropically distributed, with Crumbs2 exhibited a stronger anisotropy being almost absent from some junctions, and Myosin2B observed on all junctions but at varying levels (Figure 4B and Figure 4-figure supplement 1B).

**Figure 4:**
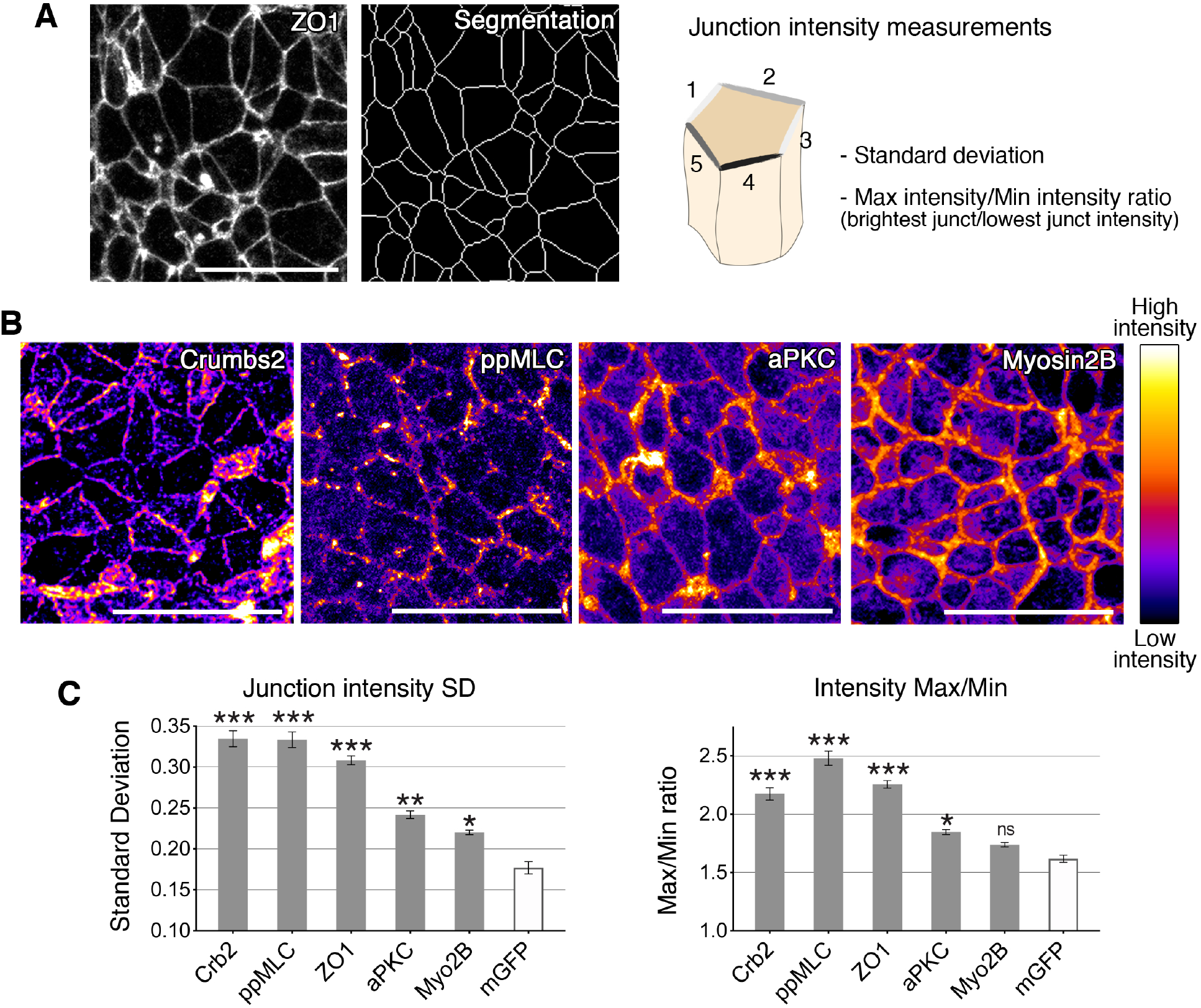
Anisotropic distribution of apical components in epiblast cells at the primitive streak. **(A)** ZO-1 apical membrane localization and corresponding skeleton of membrane segmentation used to quantify junctional intensities and anisotropy parameters as schematized. The average intensities of proteins associated with the junctions of individual cells are used to calculate the standard deviation between junctions, and the difference between junctions with the maximum and minimum intensities. **(B)** Lookup table (LUT) of immunostaining of Crumbs2, di-phosporylated Myosin Light Chain (ppMLC), aPKC and Myosin2B. These proteins show different distributions at apical junctions. Myosin2B and aPKC are present on almost every junction and seem to have a low anisotropy distribution, whereas ppMLC and Crumbs2 are not present on every junction and show a more anisotropic distribution. **(C)** Plots showing the anisotropy parameters, standard deviation and max/min. Proteins such as Crumbs2, ZO-1 and ppMLC show elevated parameters corresponding to a high level of anisotropy, whereas aPKC and Myosin2B show lower anisotropy. The membrane-GFP (recombined mTmG mouse reporter line) is used as control showing homogeneous distribution across all junctions and low anisotropy. (a range of 407-4135 junctions were analyzed in 3-5 embryos for each protein. Statistical test compare each proteins to mGFP). Pr: proximal, D: distal, R: right, L: left, PS: primitive streak. **P<0.01, ***P<0.001 (one-way ANOVA test). Error bars represent s.e.m. Scale bars, 20μm. **Figure supplement 1**: Complementary accumulation of Crumbs2 and Myosin2B, and anisotropy of apical components at the primitive streak. **Figure supplement 2**: Apical proteins show anisotropy accumulation but no planar polarity

We next investigated the localization of other apical proteins, including the actomyosin components, mono- and di-phosphorylated Myosin Light Chain (pMLC and ppMLC), Rock1, a Rho-associated kinase that phosphorylates and activates myosin light chain, and F-actin. We analyzed the localization of the Crumbs complex proteins PatJ, and aPKC, a regulator of non-muscle Myosin2 contexts (Biehler et al., 2021; Petrov et al., 2017; Roper, 2012; Sidor et al., 2020). Membrane segmentation allowed us to systematically quantify several parameters, including apical surface area, cell elongation, and the intensity of junctional protein accumulation (Figure 4A, Figure 4-figure supplement 1C). Two parameters were calculated based on junctional protein intensity and used to evaluate the level of anisotropy: the standard deviation of the junctional intensity measurements cells (SD), and the ratio of the maximum to minimum junctional intensity values (Max/Min) for each cell (Figure 4A). The membrane-GFP reporter served as a control for homogeneous membrane fluorescence localization with low anisotropy and the cohort of proteins analyzed exhibited different degrees of anisotropy (Figure 4B,C and Figure 4-figure supplement 1D,E). Some proteins such as Crumbs2, ppMLC, ROCK1 and ZO-1 showed a high level of anisotropy (Figure 4B,C and Figure 4-figure supplement 1D,E), though each exhibited a distinct pattern. ZO-1 was observed on all junctions, with variable intensity. Crumbs2 was present on some junctions but absent on others. ppMLC and ROCK1 also exhibited accumulation on some junctions, but low accumulation on others. Other proteins, including PatJ and F-actin, displayed distinct patterns but intermediate anisotropy. Finally some proteins, such as pMLC, Myosin2B and aPKC showed less anisotropy with lower SD and Max/Min values (Figure 4B,C and Figure 4-figure supplement 1D,E). These data allowed us to classify proteins based on their distribution and level of anisotropy. Staining observed at earlier (E7) and later stages (E8) did not show visible differences (data not shown) (Ramkumar et al., 2016).

The anisotropic membrane accumulation of proteins is relevant with respect to the planar cell polarity (PCP) of epithelial cells. The apical proteins we analyzed did not show a visible planar polarized distribution, which we sought to verify quantitatively.

We compared the planar polarity index calculated by Tissue Analyzer (amplitude and orientation of polarity) and mean junctional intensity in relation to junction orientation (Figure 4-figure supplement 2) (two ways to assess planar polarity) of apical proteins analyzed to Celsr1, a protein implicated in PCP in mouse and chick (Curtin et al., 2003; Mahaffey et al., 2013; Nishimura et al., 2012). Celsr1 exhibited a planar polarized localization and tended to accumulate on junctions oriented on left-right axis perpendicular to the proximo-distal axis, but by contrast, Myosin2B, Crumbs2, ppMLC, ZO-1 and aPKC, showed no specific axis of planar polarity.

### Reciprocal distribution of apical proteins at apical junctions

We next assessed the colocalization and correlation of the different apical proteins at junctions. Segmentation and intensity values, which served as a readout of concentration, were used to systematically compare mean junctional protein intensities and correlate localization. Some proteins such as ppMLC (di-phosphorylated Myosin Light Chain) and Rock1 showed similar patterns with comparable levels of anisotropy, and appeared to accumulate on the same junctions (Figure 5A). Co-staining revealed a broader accumulation for ppMLC. However, where Rock1 was seen to accumulate, ppMLC was always present, demonstrating strong colocalization (Figure 5A,5C), close to that observed with a positive control staining for ppMLC with two distinct secondary antibodies (Figure 5C and Figure 5-figure supplement 1C,E). F-actin accumulated on most junctions, exhibiting a distinct distribution from ppMLC, whereas F-actin and ppMMLC exhibited a high level of colocalization (Figure 5-figure supplement 1A), with high correlation of junctional intensities (Figure 5C and Figure 5-figure supplement 1A,E). Comparison of F-actin and Rock1 revealed a similar high correlation (Figure 5-figure supplement 1D,E). Thus, the correlated distributions of ppMLC, Rock1 and F-actin proteins suggests they could function at the same junctions. Despite exhibiting anisotropy, ZO-1 and aPKC were observed to be present on all junctions (Figure 4A,B and S3), with a high level of correlation, similar to a positive control (Figure 5-figure supplement 1D,E), demonstrating their accumulation on the same junctions.

**Figure 5:**
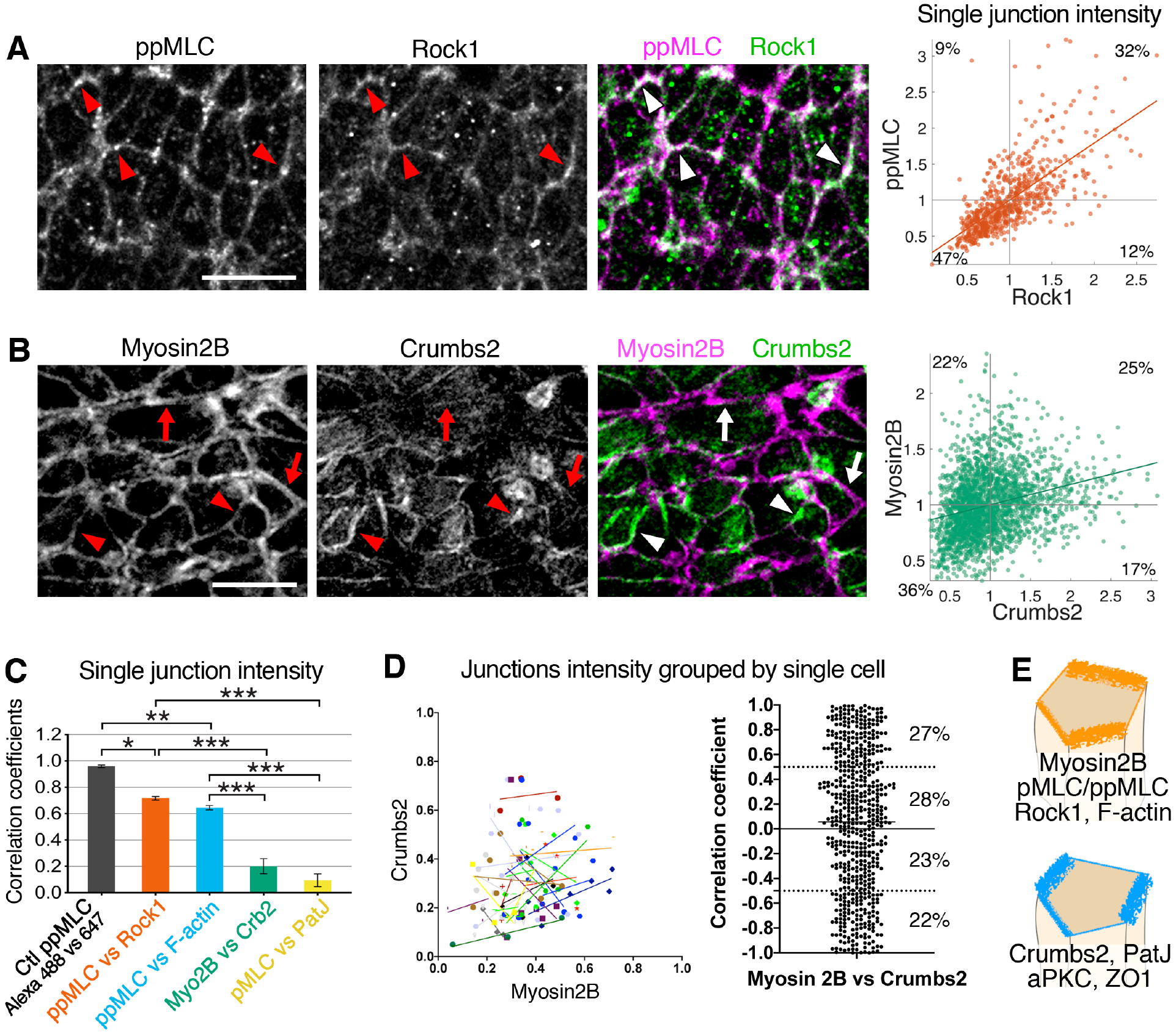
Complementarity accumulation of apical components at apical junctions. **(A)** Co-immunostaining of ppMLC and Rock1 reveals a tendency to accumulate on the same junctions. Note colocalization of fluorescent signal on most junctions (arrowheads). Scatter plot on the right showing elevated intensity correlation of the two proteins. Each point represented the mean intensity at a single junction. **(B)** Co-immunostaining of Myosin2B and Crumbs2 reveals that they tend to accumulate on different junctions. Note junctions where Myosin accumulation predominates (arrows), and junctions where Crumbs2 predominates (arrowheads). Corresponding scatter plot shows a low correlation. **(C)** Plot showing the correlation coefficients associated with the different scatter plots. Proteins such as ppMLC, Rock1 and F-actin exhibit a strong correlation with a high coefficient (ppMLC vs Rock1=0.72±0.01, ppMLC vs F-actin=0.65±0.02), whereas others including Myosin2B and Crumbs2, PatJ and pMLC exhibit much lower correlation with low coefficient (Myosin2B vs Crumbs2=0.20±0.06, pMLC vs PatJ=0.09±0.05). **(D)** Example of scatter plot showing the junctional intensity correlation of Crumbs2 and Myosin2B in a cohort of cells. Each symbol represents one junction, grouped by cells (color coded) with respective correlation trends, depicting a mix of cells with correlation, some with no correlation, and some with anti-correlation. Graph on the right shows correlation coefficients for a larger number of cells, showing a broad distribution, from correlation (+1) to anti-correlation (−1). % show the distribution of correlation coefficients in the different quarters. **(E)** Based on co-staining for multiple apical proteins and measurements of anisotropy and correlation, two groups of proteins can be defined with a tendency to accumulate on same junctions: actomyosin components, and apical determinants and junctional proteins. 711-2453 junctions were quantified in 3-4 embryos for each pair of proteins. *P<0.05, ***P<0.001 (one-way ANOVA test). Error bars represent s.e.m. Scale bars, 10μm. **Figure supplement 1**: Correlation of protein distributions at apical junctions. **Figure supplement 2**: Junctional correlation of proteins at the single cell level.

By contrast, when comparing other junctional proteins, we observed very low or no correlation. Crumbs2 and Myosin2B exhibited a complementary localization (Figure 5B), with minimal colocalization. Comparisons of Crumbs2-Myosin2B junctional intensities revealed a broad distribution and very low correlation (Figure 5B,C). Comparing PatJ, a Crumbs partner protein, with pMLC (phosphorylated-Myosin Light Chain) showed a similar accumulation with almost no correlation (Figure 5C and Figure 5-figure supplement 1B,E). Analysis of other protein pairs, including ZO-1-ppMLC, and aPKC-ppMLC revealed low correlation (Figure 5-figure supplement 1D,E).

To gauge the non-complementarity and assess potential anti-correlation of certain proteins, we analyzed junctional intensity correlations grouped by individual cells, rather than junctions collectively. This analysis confirmed the strong correlation of ppMLC, Rock1 and F-actin, where most cells exhibited a positive correlation trend with a high correlation coefficient (Figure 5-figure supplement 2A,B). For pairs of proteins such as Crumbs2 and Myosin2B, and PatJ and pMLC, which showed no correlation when junction quantifications were compared (Figure 5B,C and Figure 5-figure supplement 1), grouping junction intensities by individual cells revealed a broad distribution; some cells showed anti-correlation, some no correlation and some positive correlation (Figure 5-figure supplement 2C-F). This analysis illustrated a heterogeneity across the epiblast population in the vicinity of the primitive streak, at any one time, supporting our hypothesis that anisotropic accumulation patterns are dynamic.

These quantitative analyses of apical protein anisotropy patterns combined with the correlation of distributions, allowed us to define two groups of proteins that tend to accumulate on different junctions. Cytoskeletal proteins including Myosin2, phosphorylated MLC, ROCK1 and F-actin predominantly accumulate on the same junctions, while apical components and junctional proteins such Crumbs2, PatJ, aPKC, ZO-1 will co-localized on other junctions (Figure 5D). This anisotropic distribution of the actomyosin network and Crumbs-complex associated proteins is associated with dynamic apical constriction.

### Cellular defects in *Crumbs2^−/−^* mutant embryos at the primitive streak

Crumbs2 is an important regulator of the mouse gastrulation EMT, where it promotes cell ingression. Epiblast cells fail to ingress and accumulate at the primitive streak in *Crb2*^−/−^ embryos (Ramkumar et al., 2016). By quantifying the apical surface of posterior epiblast cells at E8.0, we observed increased cell density and cells with smaller apical surfaces around the primitive streak of mutants, as compared to littermate controls (Figure 6,7-figure supplement 1A). This cell crowding was not observed half a day earlier at E7.5 (Figure 6,7-figure supplement 1B) (Ramkumar et al., 2016) when mutant embryos were morphologically indistinguishable from wild-type stage-matched counterparts.

Morphological defects (apical surface areas and surface elongation) in cells at the primitive streak were first detected at E8.0 with cells having smaller and rounder apical surfaces (Figure 6A-E and Figure 6,7-figure supplement 1B) resulting from their failure to ingress, and consequent accumulation in the epiblast.

**Figure 6:**
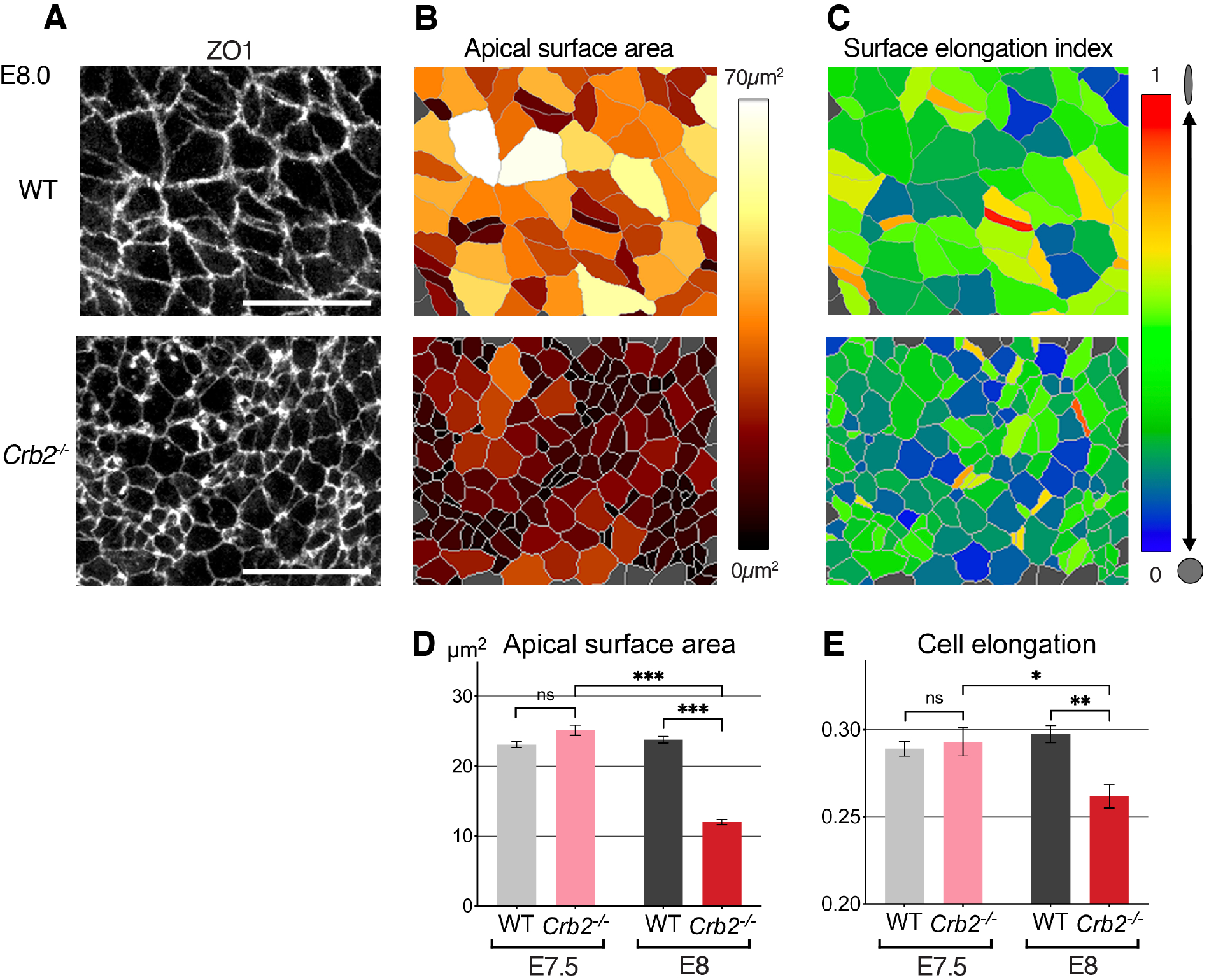
Cellular defects in Crumbs2 mutant embryos at the primitive streak. **(A)** ZO-1 immunostaining showing the apical surface of WT and *Crumbs2* mutant embryos at E7.5 and E8.0. **(B)** Segmentation and color code of apical surface areas reveals smaller surfaces in *Crumbs2* mutants at E8.0. **(C)** Color code of cell surface elongation index calculated by Tissue Analyzer (regardless of their axis) reveals rounder surfaces in *Crumbs2* mutants at E8.0. **(D)** Graph of quantifications showing significantly smaller apical surfaces in *Crumbs2* mutants at E8.0 compared to E7.5, and to WT embryos. **(E)** Graph showing significantly less elongated apical surfaces in *Crumbs2* mutants at E8.0 compared to E7.5, and to WT embryos. 334-1187 cells from 3-5 embryos were quantified for each genotype and stage. ***P<0.001 (Unpaired bilateral Mann-Whitney test). Error bars represent s.e.m. Scale bars, 20μm.

### Molecular defects in *Crumbs2^−/−^* mutant embryos precede cellular defects

To identify molecular defects associated with and preceding the ingression defect we analyzed the distribution of actomyosin components in *Crb2*^−/−^ embryos. Although ZO1 distribution appeared unchanged in the mutants, suggesting the integrity of junctions was not altered, we observed reduced levels of Myosin2B and ppMLC (the fully activated form of Myosin Light Chain) in mutants at E7.5 (Figure 7A,B) and Figure 6,7-figure supplement 1C). We also observed more diffuse and reduced cortical actin filaments at the apical junctions of epiblast cells in mutants, whereas the medial pool of actin appeared slightly increased (Figure 7A,B). Thus, actomyosin apical accumulation and activity were perturbed without alteration of junctional integrity in *Crb2*^−/−^ embryos at E7.5, preceding morphological defects.

**Figure 7:**
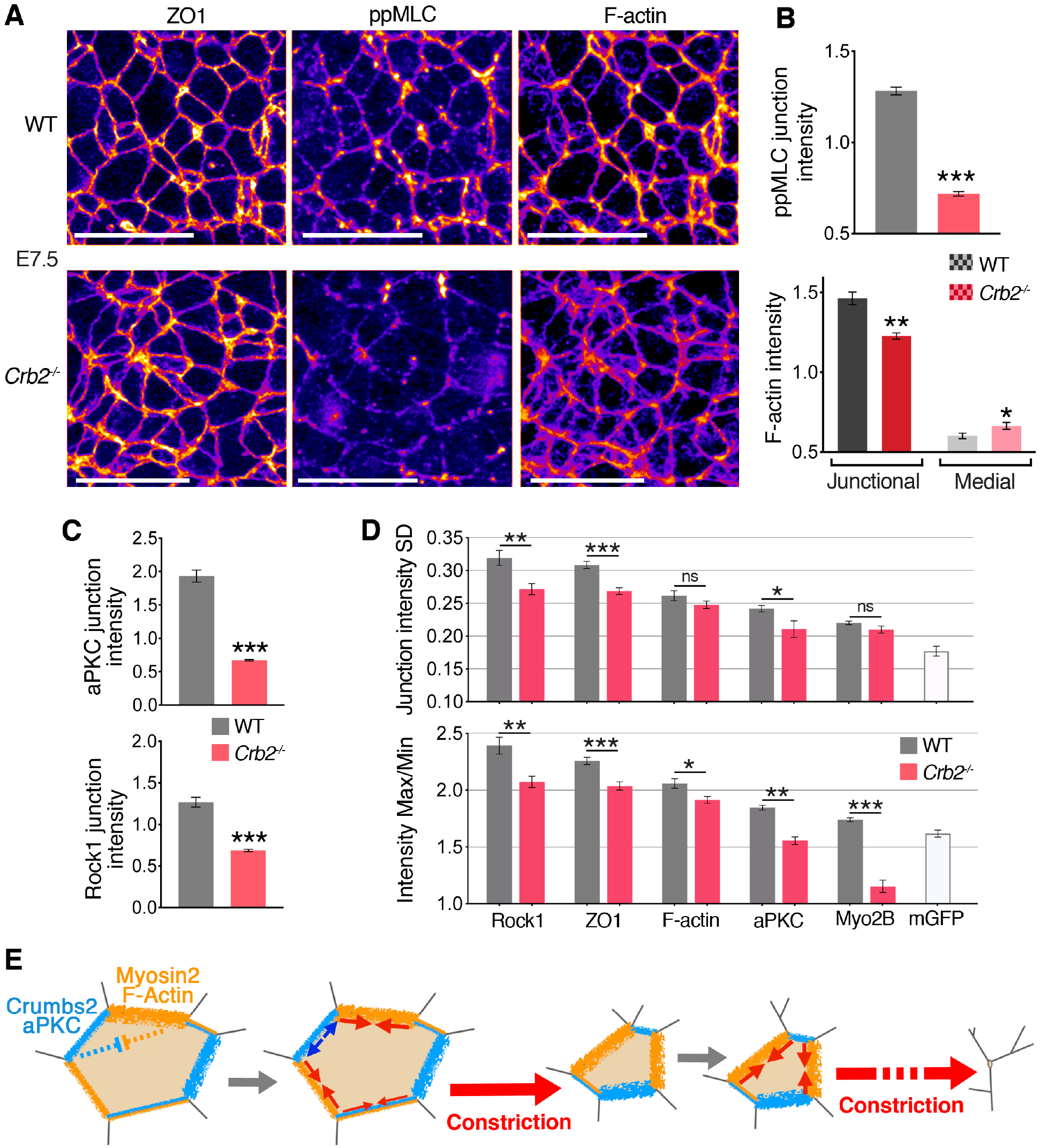
Molecular defects in *Crumbs2* mutants preceding visible cellular defects. **(A)** Co-immunostaining for ZO-1, ppMLC and F-actin in WT and *Crumbs2* mutants at E7.5 reveal reduced ppMLC and F-actin junctional intensities. **(B)** Plots showing intensity measurements of junctional ppMLC and F-actin. ppMLC shows significant decrease in junctional accumulation in *Crumbs2* mutants. F-actin shows lower junctional accumulation and increased diffuse medial intensity in *Crumbs2* mutants. **(C)** Graphs of junctional intensity measurements showing significantly decreased aPKC and Rock1 intensity in *Crumbs2* mutants compared to WT. **(D)** Anisotropy parameters, Standard Deviation and Max/Min showing perturbed junctional accumulation anisotropy in *Crumbs2* mutants compared to WT embryos. ZO-1, F-actin, Myosin2, aPKC and Rock1 show reduced anisotropy in *Crumbs2* mutants. **(E)** Working model. Actomyosin network and Crumbs2 complex show anisotropic and complementary distribution at apical junctions of epiblast cells at the primitive streak. These distributions are likely dynamic through time and heterogenous among cells, and could potentially regulate each other distribution. Junctions asynchronously shrink and allow ratchet-like constriction of apical surfaces, facilitating cell constriction and ingression without creating increased tension within the tissue. 122-501 cells in 3-4 embryos were quantified for each genotype and proteins. *P<0.05, **<0.01, ***P<0.001 (B,C: Unpaired bilateral Mann-Whitney test; D: one-way ANOVA test). Error bars represent s.e.m. Scale bars, 20μm. **Figure supplement 1**: Cellular and molecular defects in *Crumbs2* mutant embryos.

We next focused on aPKC and Rock1, implicated in the regulation of the actomyosin network by Crumbs in Drosophila and cultured cells (Biehler et al., 2021; Ishiuchi and Takeichi, 2011; Roper, 2012; Sidor et al., 2020). By quantifying the intensity of aPKC and Rock1 at junctions, we observed a 3-fold reduction of aPKC intensity at apical junctions and a more diffuse signal in the cytoplasm, and we observed a slight reduction of junctional Rock1 (Figure 7C and Figure 6,7-figure supplement 1D). These observations suggest a network with aPKC and Rock1 regulating actomyosin downstream of Crumbs2. By quantifying the two parameters used to assess anisotropic distributions at apical junctions, we observed that in *Crb2*^−/−^ embryos the SD and Max/Min of junction intensity of Myosin2, F-actin ZO-1, aPKC and Rock1 was reduced (Figure 7D), revealing reduced anisotropy and more homogenous distribution of these proteins.

## DISCUSSION

To develop a dynamic mechanistic understanding of the mouse gastrulation EMT at cellular resolution and tissue-scale we 3D time-lapse imaged *ex utero* cultured embryos. We used membrane tagged reporters and quantified the spatiotemporal features of apical cell surfaces, junctions and junctional composition of cells in the vicinity of the primitive streak. Our analyses reveal that after converging at the primitive streak, epiblast cells undergo EMT, either individually or in small groups of cells, in what appear to be stochastic events. Cells constrict their apical surfaces in a ratchet-like manner during ingression from the epiblast epithelium. Constriction is associated with the anisotropic apical distribution of actomyosin network and Crumbs2 complex proteins.

Apical constriction is an evolutionarily conserved behavior associated with gastrulation in invertebrates such as Drosophila, nematodes and sea urchins, as well as in vertebrates and amniotes, such as Xenopus and chick (Christodoulou and Skourides, 2015; Lecuit and Lenne, 2007; Martin and Goldstein, 2014; Martin et al., 2009; Rozbicki et al., 2015). We observed fluctuations in the size and shape of the apical surface of epiblast cells as they converged towards the streak. In the vicinity of the streak, and as cells ingressed, their apical surfaces predominantly constricted, in a step-wise ratchet-like manner.

Pulsed ratchet-like apical constrictions have been observed in Drosophila gastrulation, where cells at the ventral furrow constrict in a short period of time to drive tissue invagination, followed by en-masse cell delamination (Martin et al., 2009; Mason et al., 2013). Drosophila neuroblast delamination also exhibits a ratchet-like apical constriction (An et al., 2017; Simoes et al., 2017). Despite these similarities, mesoderm formation process during gastrulation of Drosophila and mouse cannot be directly compared as they exhibit different time-scales, mechanisms and spatial parameters. In Drosophila the entire ventral furrow constricts in less than 10min, triggering a tissue-wide invagination without cell ingression, thereafter cells lose their epithelial properties and form a mesenchymal mesodermal tissue. By contrast, in the mouse embryo epiblast cells at the primitive undergo constriction and ingression in a more isolated fashion over the course of almost 2 days. Our time-lapse imaging revealed a ratchet-like apical cell constriction operating in the mouse gastrulation EMT, where cells predominantly ingress as isolated cells or small cluster, versus the en-masse constriction and invagination taking place in Drosophila.

We further demonstrated that this apical constriction is associated with the asynchronous contraction of junctions between neighboring cells, potentially regulated by the apical actomyosin network. During Drosophila gastrulation, an apico-medial network of actomyosin pulls apical cell junctions to simultaneously reduce their length and allow the constriction of apical surface of cells at the ventral furrow (Coravos and Martin, 2016; Martin et al., 2009; Mason et al., 2013). Drosophila neuroblasts exhibit polarized apical constriction driven by both planar polarized junctional and medial actomyosin networks, predominantly leading to a loss of antero-posterior junctions (Simoes et al., 2017). In the mouse gastrula, we found that cells undergoing EMT contract and lose cell-cell junctions in an asynchronous manner without apparent planar polarity. Our analyses of the junctional distribution of Myosin2 supports its potential involvement in regulating constriction during ingression, but does not exclude an additional role played by a medial pool. Pulsed actomyosin activity has been documented in various processes involving cell shape changes, including epithelial apical constriction (Coravos et al., 2017; Heer and Martin, 2017; Martin and Goldstein, 2014). The pulsed constriction we observe in mouse gastrulation indicates conservation of this actomyosin behavior, as a general mechanism that integrates assembly and disassembly of the network and rearrangement and recruitment of Myosin and Actin molecules. Furthermore, pulsed contractility could allow a substantial but transient constriction of epiblast apical surfaces during ingression without creating deleterious tension which could impact the integrity of the tissue.

Since large numbers of cells undergo EMT and leave the epiblast epithelium at any given time, simultaneous constriction and ingression of multiple neighboring cells could exert extreme tension and deformation in the tissue. Our observation of a stochastic and asynchronous constriction of neighboring cells provides a parsimonious explanation for maintaining epithelial integrity, and a continuation of ingression events at the primitive streak and tail bud for almost 2 days throughout gastrulation, in comparison to Drosophila gastrulation where all ventral furrow cells constrict in less than 10min. Cell division taking place on the basal side of an epiblast layer, has been reported at the primitive streak where it has been proposed to facilitate cell ingression (Mathiah et al., 2020). Apical constriction and basal cell division could therefore work in concert to promote the correct number of cell ingression events.

Myosin2 is a driver of membrane movement and apical constriction in various contexts (An et al., 2017; Chung et al., 2017; Marston et al., 2016; Martin and Goldstein, 2014; Martin et al., 2009; Mason et al., 2013; Nishimura et al., 2012; Roh-Johnson et al., 2012; Samarage et al., 2015; Simoes et al., 2017). Crumbs2 is involved in the EMT process during mouse gastrulation and exhibits a complementary localization to Myosin2 (Ramkumar et al., 2016) (our data). Anisotropic membrane localization of Crumbs has been implicated in Drosophila salivary gland development, where it controls the localization of actomyosin cables at the placode border, necessary for apical constriction and tissue invagination (Roper, 2012). Recently, Crumbs has also been implicated in apical constriction during neuroblast cell ingression in Drosophila (Simoes et al., 2022). We observed the anisotropic and complementary localization of Crumbs2 and Myosin2, and have identified other proteins exhibiting anisotropic distributions. Although Myosin2B, F-actin, aPKC and ZO-1 are present on all apical junctions, they exhibit a differential anisotropic accumulation between the junctions of any one cell. Other proteins such as ppMLC and the myosin kinase Rock1, as well as Crumbs2 and its partner PatJ, exhibited a higher level of anisotropy. In the Drosophila salivary gland placode border, Crumbs and Myosin2 show planar polarized localization with Crumbs being absent from cell junctions that contact the border of the placode, allowing the accumulation of myosin II at the placode border and formation of a supracellular actomyosin cable (Roper, 2012). By contrast, our results in mouse show that Crumbs2, Myosin2B and several other proteins do not exhibit planar polarity, but what appears to be a disordered anisotropy. Our studies reveal a patterned distribution in which actomyosin regulators and Crumbs2, aPKC and ZO-1 accumulate on subsets of cell junctions and display complementary distributions. We propose that these anisotropic patterns are dynamic and control the asynchronous junction contraction associated with apical constriction.

Crumbs2 acts through aPKC and Rock to regulate Myosin2 localization in the *Drosophila* salivary gland (Roper, 2012), and aPKC has been shown to regulate Myosin2 junctional localization and activity both positively and negatively in different context, directly or through other proteins (Biehler et al., 2021; Ishiuchi and Takeichi, 2011; Petrov et al., 2017). The regulation of Myosin2 by Crumbs and aPKC through Rock in *Drosophila* salivary gland development is complex (Sidor et al., 2020), and despite Rock membrane association being tightly regulated by Crumbs across the tissue, regionally distinct behaviors have been observed. At placode borders, cells exhibit a polarized accumulation of Crumbs/aPKC and Rock to control actomyosin cable formation, whereas in the center, where cells constrict during the tissue invagination, both Crumbs and Rock are isotropically present at junctions (Chung et al., 2017; Sidor et al., 2020). Our results demonstrate that during mouse gastrulation Crumbs2 controls the apical accumulation and activity of actomyosin, which could account for the defects in apical constriction and ingression in Crumbs2 mutants. We observed Crumbs2 as required for the correct accumulation and distribution of aPKC, Myosin2, Rock1 and F-actin at apical junctions, without alteration of junctional integrity. Crumbs2 along with aPKC could control the anisotropic accumulation of actomyosin, triggering asynchronous junctional reduction resulting in apical constriction during ingression (Figure 7E).

In ingressing mouse epiblast cells Crumbs2 together with aPKC and Rock1 could represent a complex network implicating other proteins, required for Myosin2 localization and activity at the apical membrane, but also potentially regulating the anisotropic accumulation of the actomyosin network locally at junctions, facilitating correct apical constriction during ingression (Figure 7E). Our work reveals that pulsed ratchet-like constriction behaviors observed in different species and distinct contexts are conserved in the mouse gastrulation EMT.

## METHODS

### Experimental animals

FVB or mixed background mice were used for immunofluorescence staining. Most other experiments, incorporating genetically engineered mouse models, were carried out in a mixed background. The *Gt(ROSA)26Sor^tm4(ACTB-tdTomato,-EGFP)Luo^*/<u>J</u> (Muzumdar et al., 2007) (referred to throughout as mT/mG) line was obtained from the Jackson Laboratory and crossed to Sox2-Cre (Hayashi et al., 2002) mice to generate a membrane-GFP reporter. The ZO-1-GFP protein fusion knock-in mouse reporter a gift from Terry Lechler (Duke University Medical Center, NC, USA) (Foote et al., 2013; Huebner et al., 2014). *Crumbs2^+/−^* animals were generated from animals carrying a conditional allele of Crumbs2 via germline deletion (Alves et al., 2013). Animals were housed and bred in accordance with institutional IACUC guidelines. The MSKCC IACUC approved all experiments.

### Embryo dissection, immunostaining and antibodies

For the majority of immunostaining experiments, embryonic day E7.5 and E8.0 embryos were dissected in cold PBS and fixed for 1 hour in 4% PFA, with the exception that 8% PFA was used for detection of phosphorylated myosin. Whole-mount embryos were blocked overnight in blocking buffer (PBS; 0,1%Triton; 3%BSA), incubated overnight at 4°C with primary antibodies, washed in blocking buffer and incubated overnight at 4°C with species-specific secondary antibodies. For staining of F-actin embryos were incubated with Phalloidin-FITC/TRITC (Invitrogen).

The following antibodies were used: rabbit anti-Myosin Heavy Chain 2B (1/400, Biolegend 909902), mouse anti-Myosin Heavy Chain 2B (1/50, DSHB CMII 23), rabbit anti-pMLC (1/200, Cell Signaling Ser19 3671), mouse anti-pMLC (1/200, Cell Signaling Ser19 3675), rabbit anti-ppMLC (1/200, Cell Signaling Thr18/Ser19 3674), rat anti-ZO-1 (1/50, DSHB R26.4C), mouse anti-ZO-1 (1/200, Invitrogen 33-9100), rabbit anti-ZO-1 (1/200, Zymed 61-7300), mouse anti-aPKCλ (1/100, BD bioscience 610207), chick anti-GFP (1/400, Abcam 13970). Rabbit anti-Crumbs2 was a gift from Jane McGlade (Hospital for Sick Children, Toronto, Canada) and was used at 1/50 (Laprise 2006). Rabbit anti-PatJ was a gift from Andre Le Bivic (Developmental Biology Institute of Marseille, France) and was usedc at 1/200 (Lemmers 2002). Rat anti-Rock1 was a gift from Masatoshi Takeichi (RIKEN Center for Developmental Biology, Kobe, Japan) and used at 1/100 (Nishimura and Takeichi 2008). Guinea Pig anti-Celsr1 was a gift from Danelle Devenport (Princeton University, Princeton, USA) and was used at 1/500 (Devenport and Fuchs 2008). Fluorescent secondary antibodies Alexa 488, 568 and 647 from Life Technologies and ThermoFisher were used at 1/500.

### Fixed embryo epiblast mounting and imaging

Following staining, embryos were microdissected to retain the posterior side containing the primitive streak and mounted on glass slide and coverslipped in Fluoromount-G, with the cavity and apical surface of the epiblast facing the coverslip. Layers of tape were placed on both extremity of the slide to create a slight volume and to avoid extensive compression and flattening of the tissue. The position of the primitive streak was determined to coincide with the posterior midline running from the allantois to the node (two morphologically distinguishable structures), and streak axis was oriented so that the proximo-distal axis was oriented along the vertical axis of the image. Most immunostaining revealed a high concentration of puncta also allowing us to identify the location of the primitive streak after imaging. Embryos were imaged on a Leica SP8 or a Zeiss LSM880 laser point scanning confocal using x40 oil immersion objectives, and images were acquired as Z-stacks with 1μm step.

### Segmentation and quantification of fixed embryo staining

Confocal images were analyzed using ImageJ. The primitive streak region was located and a maximal intensity projection of a minimal thickness (5-10μm) was generated to observe the apical surface of cells in a selected region of interest at the primitive streak. ZO-1 localization was generally used as an apical junction marker for segmentation.

Segmentation was performed using the Tissue Analyzer software (Aigouy et al., 2010; Aigouy et al., 2016), an ImageJ plugin which uses a watershed algorithm to segment the cell cortex. Briefly, a first membrane segmentation was automatically performed, and then verified and corrected manually. Cells and junctions were tracked and assigned IDs. Most cell parameters used were calculated by the software, including; apical surface area, cell elongation magnitude and orientation, number of junctions per cell, and signal intensities along junctions. In brief, cell surface elongation is characterized by an axis and magnitude, and can be represented by a symmetric tensor at the centroid of a cell (images with red nematic bars). Calculation integrates cell area and shape so that cells of similar shape with different sizes are assigned the same value of cell elongation (see Aigouy et al. 2016 for details equations (Aigouy et al., 2016)).

Color-code representations of apical surface area, surface elongation, and junction number were generated in Tissue Analyzer. Surface elongation index (magnitude) calculated by the software, was represented as an index from 0 (round) to 1 (line). Surface elongation and planar polarity of proteins were characterized by an axis and magnitude, represented by a symmetric tensor at the centroid of a cell (images with red nematic bars).

Intensity measurements of immunostaining are presented for every data as the mean intensity along single junctions and normalized to the global mean junctional intensity of corresponding embryos. Anisotropy parameters were calculated from junctional intensities measured within Tissue Analyzer. Intensities were measured along 3 pixel-wide lines (~500nm), measurements were extracted from a database created by the software and used externally for calculations. For each single cell analyzed, the mean junction intensity was used to calculate per cell, the standard deviation between the intensity of all junctions, and the junction with the maximal intensity divided by the junction with the minimal intensity (Max/Min ratio). These two parameters were used to evaluate the anisotropic accumulation of proteins at apical junctions. Double immunostainings were used to assess correlative accumulation of proteins, and mean junction intensities were used to create scatter plots to evaluate the dispersion of the intensity across the populations, and their correlation coefficients. Generally, measurements taken of junctions from different embryos were pooled, except for a few examples in which junctions were grouped by individual cells to evaluate the range of anisotropy in different cells.

For planar polarity of protein localization at apical junctions, Tissue Analyzer calculates a planar polarity characterized by an axis and magnitude, and can be represented by a symmetric tensor at the centroid of a cell (images with red nematic bars). Cell junctions fluorescent intensity as well as the angles of junction and the geometry of the cell were used to define the magnitude of polarity as well as the angle (see Aigouy et al. 2016 for details equations (Aigouy et al., 2016)). For protein accumulation based on junctional orientation, average junctional intensities were plotted according to junctional orientation angles which were classified in bins of 15°.

For protein quantifications in *Crb2*^−/−^ mutants and comparison with wild-type (WT) embryos, 3-4 embryos of each genotype recovered from 2-3 litters where processed, immunostained and imaged in parallel using the same parameters, then segmented and quantified as described above. Measurements are presented as the mean intensity along single junctions and normalized to the global mean junctional intensity of the pool cohort of embryos of both genotypes for each proteins. For F-Actin quantification, junctional values represent the intensity measured at the peripheral cortex and the medial values the intensity measured in the medial/cytoplasmic area of the same cells.

### Time-lapse imaging

For time lapse imaging, mid- to late-streak stage embryos (E7.5) were dissected with their yolk sac and ectoplacental cone intact in dissection medium (CO_2_ independent DMEM, 10%FBS) at 37°C. Embryos were stabilized in a depression created in a layer of collagen matrix and imaged in glass-bottom 35mm MatTek dishes. They were placed with their posterior side facing the objective and culture in 50% rat serum/50% DMEM media in an incubation chamber at 37°C and 5% CO_2_. Embryos were imaged for a total of 3-6h with Z-stacks of 1μm step acquired every 5min (minimum time-scale possible to keep embryo and fluorescent signal integrity) on a Leica-SP8 laser point scanning confocal or a Nikon A1RHD25 high speed resonant scanning confocal with 40x objectives. Imaging usually focused on regions of interest in the proximo-distal middle of the primitive steak where the epiblast is the least curved and optimal for image data acquisition and analysis.

### Dynamic images analysis and quantification

Membrane-GFP from *mT/mG* embryos was used to observed apical constriction associated with cell ingression at the primitive streak. For all other imaging and quantification, a ZO-1-GFP Knock-in reporter was used as it generally gave the best signal-to-noise enabling observation of apical surfaces of epiblast cells. Analyses were performed in ImageJ and Tissue Analyzer on maximal projections of Z-stacks. Cell tracking in the plane of the epithelium was performed manually using the Manual Tracking plugin in ImageJ. To quantify ingression events, small areas of interest in the vicinity of the primitive streak, corresponding to a ~40μm region along the predicted midline, were analyzed. All cells were tracked for at least one hour, and constriction and ingression events were identified to calculate the ratio of ingression events per hour. Ingression events were considered isolated when the constriction and disappearance of a cell occurred within 30min or more of the ingression of adjacent neighboring cells, and clustered when occurring within a 30min window of the ingression of adjacent neighboring cells (2-4 cells in different clustered events).

For dynamic analysis of apical surfaces, small regions of interest at the primitive streak were used for segmentation with Tissue analyzer. A first segmentation was automatically generated by the software, then each time-point manually corrected. Cell parameters, including apical surface area, surface elongation index and orientation and number of edges, were extracted from the segmentation database, and graphs were plotted using GraphPad Prism.

The beginning of cell constriction associated with an ingression event was defined as the point when cells started to reduce their apical surface area after a period of oscillation, and so continually until they completely lost their apical domain. The start of constriction was defined as t0 when the apical surface area, rate of change of surface area and other parameters were plotted against time. Changes in apical surface area (constriction rate) was defined as the inverse value of the derivative of apical surface area: Δarea(t)=area(t-1) – area(t). Oscillations of surface area in epiblast cells away from the primitive streak, and before the initiation of ingression were compared with the more dramatic changes in surface area during the constriction period. During ingression, a constriction pulse was defined as an event in which the contraction rate exceeded one standard deviation above the mean of the contraction rate of non-ingressing cells (above 0.7 μm^2^/min). The number of pulses, occurrence of pulse and pulse magnitude were quantified manually for each constriction period of each cell and each pulse. The calculated apical area rate of change was integrated in the Tissue Analyzer database, and used to generate color-coded movie representations of surface area, area rate of changes, cell elongation, number of junctions and junction length. As for fixed embryo analysis, surface elongation index (magnitude) calculated by the software, is represented as an index from 0 (round) to 1 (line). Surface elongation is characterized by an axis and magnitude, and is represented by a symmetric tensor at the centroid of a cell (red nematic bars).

Changes in junction length (shrinkage rate) was defined similarly to cell area changes, as the inverse value of the derivative of junction length: Δlength(t)=length(t-1) – length(t). To analyze and compare junctions length changes over time, correlation of pairs of junctions were quantified and plot as correlation matrix to illustrate a single cell or correlation coefficient plot for pool of single cells. The average shrinkage rate of single junctions during consecutive cell constriction pulses and stable phases were calculated to visualize the asynchronous and differential shrinkage of junctions during single cell constriction.

### Statistics

A range of 3 to 6 embryos were analyzed for each experiment. Details of n values, means and p values can be found in Supplementary file 1. Error bars on graphs represent s.e.m. P values corresponds to an unpaired non-parametric Mann-Whitney test for comparison of pairs of conditions, or one-way ANOVA for multiple comparisons. All cells were included in the statistical analysis. Each embryo was considered a biological replicate. No randomisation or blind analysis were conducted.

### Definition of terms

#### Start of constriction

the beginning of cell constriction associated with an ingression event at the primitive streak was defined as the time point when cells started to reduce their apical surface area after a period of oscillation, and so continually until they completely lost their apical domain.

#### End of constriction

the end of constriction was defined as the time point when the constricting cells lost their apical surface and move out of the plan of the epiblast.

#### Before constriction

defines the time period before cells start their constriction at the primitive streak. This time-period shows same oscillation characteristics of the apical surface area as epiblast cells further away from the primitive streak.

#### Constriction

defines the time-period during which cells reduced their apical surface area until they completely lose their apical domain. We also use the term for the constriction rate and constriction pulse.

#### Contraction/expansion

used to define the rate of change in apical surface area through time, contraction defining the rate of reduction in area, and expansion a rate of increase in area. They are defined both as positive value and plotted as rate of change, or in some cases defined negatively and positively and plotted as constriction rate.

#### Constriction pulse

defines the time-period during the constriction when the rate of contraction of the apical surface area exceeds the threshold.

#### Stable phases

defines the time-period between constriction pulses, when the change in apical surface area shows minimal variation.

#### Pulse occurrence

defines the frequency at which constriction pulses are occurring. Pulse magnitude: defines as the maximum value of constriction rate within each pulse. Shrinkage: defines the junction reduction in length, Shrinkage rate indicates the change in junction length through time.

## Acknowledgements

AF and AKH dedicate this manuscript to the memory of our late colleague and mentor Kathryn V Anderson. Kathryn marveled at the spectacle of mammalian gastrulation and recognized the insights that genetics and imaging would bring. We thank Andre Le Bivic for the PatJ antibody, Danelle Devenport for the Celsr1 antibody, Jane McGlade for the Crumbs2 antibody and Masatoshi Takeichi for the Rock1 antibody. We thank Terry Lechler for the ZO-1-GFP animals. We thank Zhirong Bao, Alex Joyner, Jen Zallen and members of the Hadjantonakis lab for stimulating discussions and critical feedback on the work. We thank the MSKCC (Memorial Sloan Kettering Cancer Center) Molecular Cytology, and the SKI Light Microscopy Instrument Cluster and James Muller for imaging technical support. This study was supported by the NIH (R01HD094868, R01DK127821, R01HD086478, and P30CA008748). AF was supported by a postdoctoral fellowship from the MSKCC GMTEC (The Alan and Sandra Gerry Metastasis and Tumor Ecosystems Center) for part of this work.

---

**Figure 1-figure supplement 1:**
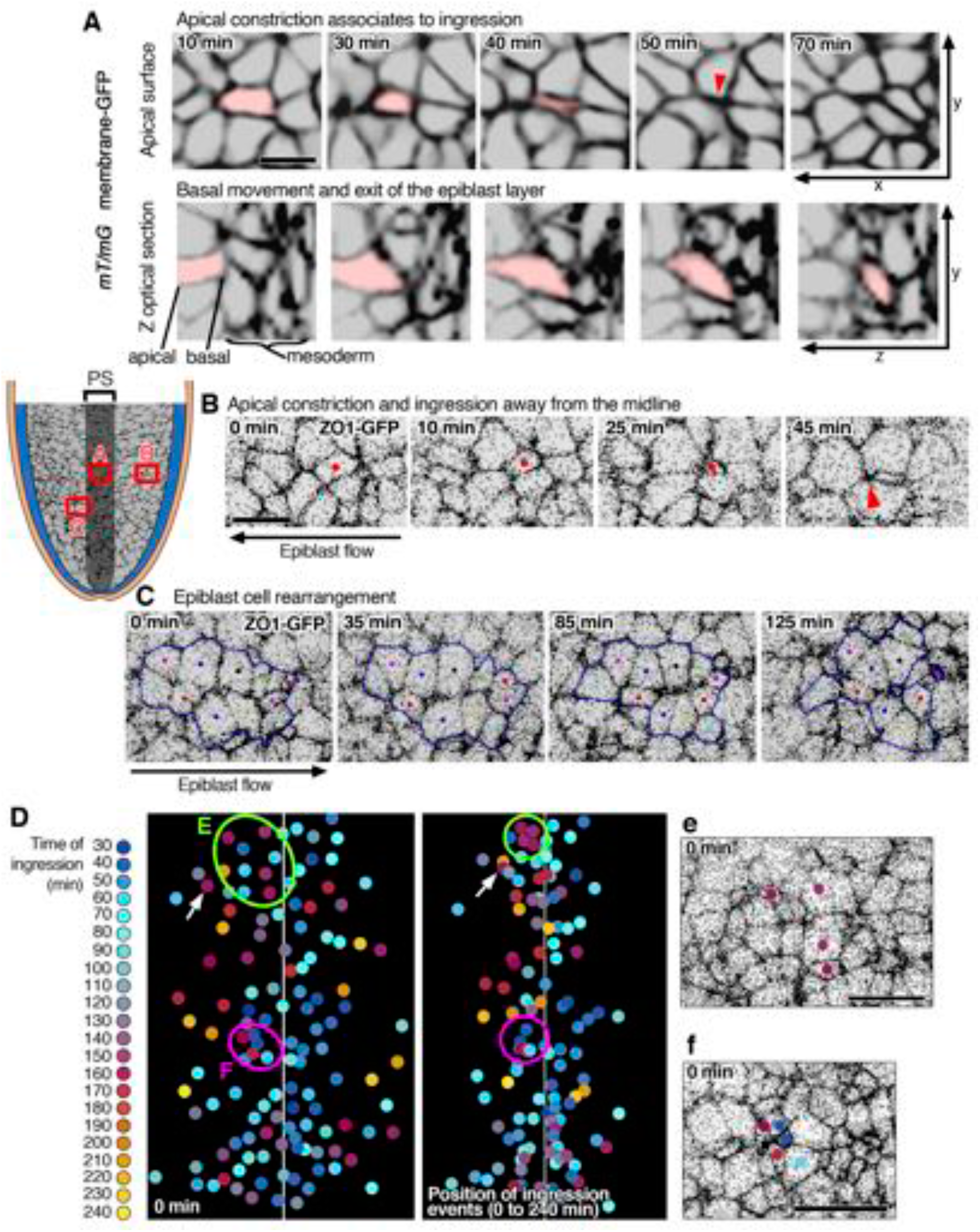
Apical constriction and ingression of epiblast cells in the vicinity of the mouse primitive streak. **(A)** Snapshots of a time-lapse of membrane-GFP in an *mTmG* embryo showing apical surface constriction in a single Z plan below the apical membrane (top row) associated to ingression out of the epiblast layer and integration to the mesoderm seen on apicobasal Z optical section (bottom row) at the primitive streak (region located on the schematic). **(B)** Example of cell constricting away from the midline, as the epiblast cells move toward the primitive streak. **(C)** Example showing cell rearrangement among a cluster of epiblast cells near the primitive streak. **(D)** Color-coded time-series and position of ingression events of tracked cells shown in Figure 1d,e. The initial position of a circled group of cells (green circle) is shown in e; cells are not all neighbors at 0min but ingress together as neighbors (after 165min). Another group of cells (purple circle) who’s initial position is shown in f; cells are neighbors at 0min, but they do not ingress at the same time. Cells are considered to constrict and ingress as isolated cells when their ingression takes place 30min or more from their neighbors. White arrow shows the ingression of an isolated cell. Scale bars A,B,C 10μm; E,F 20μm.

**Figure 2,3-figure supplement 1:**
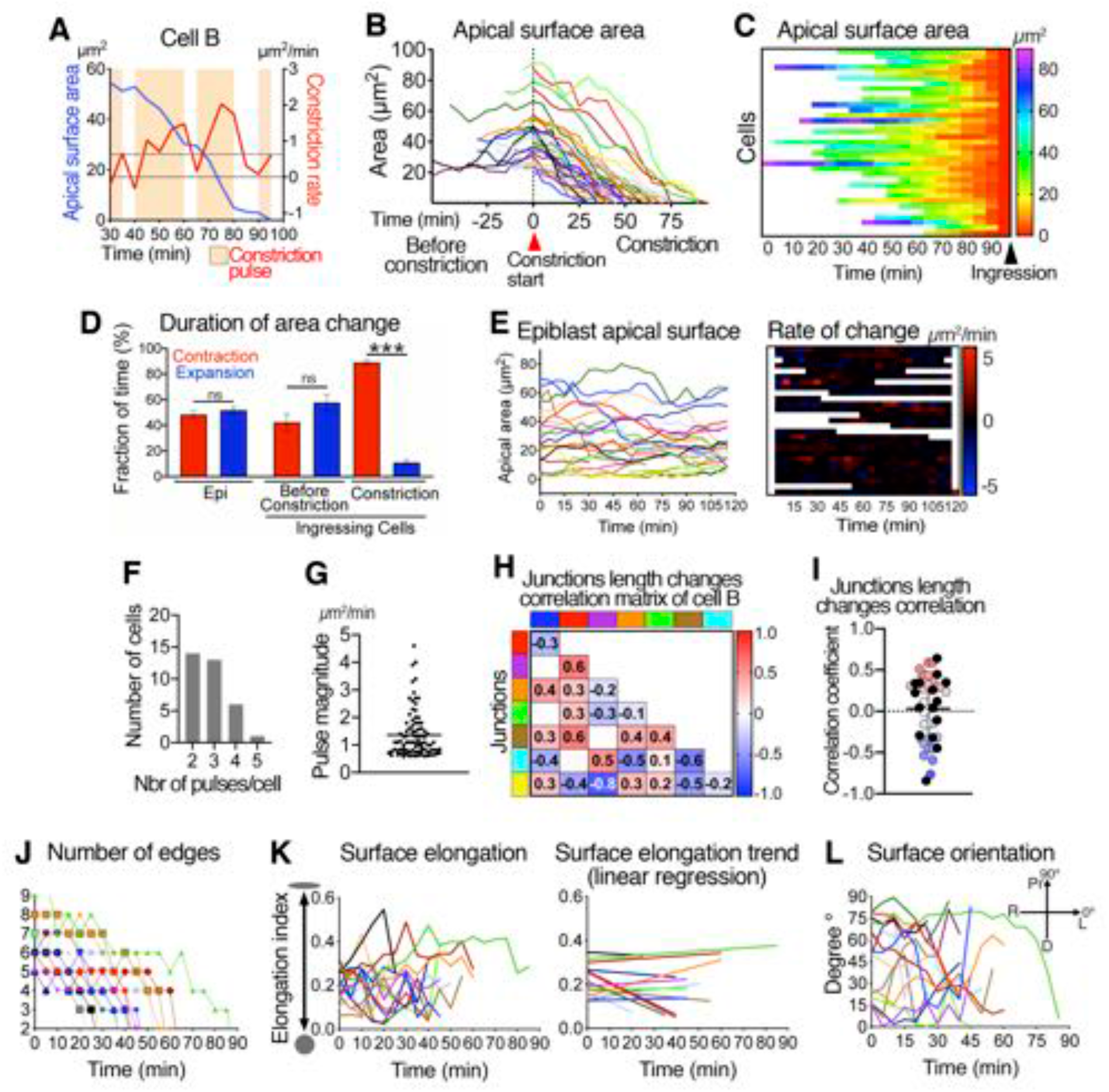
Details and features of constriction pulses and cell parameters during constriction. **(A)** Graph of apical surface area and rate of constriction of single cell B (shown in Figure 3D,E) exhibiting four pulses of constriction (beige regions). **(B)** Plot showing the apical surface area of several cells at the primitive streak during the total time of the analysis. 0min marks the beginning of the constriction period. Cells begin with a wide range of different initial apical surfaces prior to their constriction. **(C)** Heat map plot of the apical surface area during constriction showing the reduction in area and the pulses of constriction. Each row represents data for an individual cell. **(D)** Fraction of time cells spend contracting or expending their apical surfaces revealing equilibrium in lateral epiblast and at the streak before the constriction period, followed by significantly longer contraction time during constriction. **(E)** Plot showing the apical surface area and heat map color-code of rate of change of epiblast cells outside the region occupied by the primitive streak before ingression. Cells exhibit a wide variety of apical areas, and fluctuate their apical surfaces, with a low rate of contraction and expansion through time. (Corresponding measurements are shown in D and Figure 2E). **(F)** Frequency distribution graph showing the proportion of cells undergoing two to four main constriction pulses. **(G)** Dot plot showing the average magnitude of constriction pulses. **(H)** Matrix heat map showing the correlation of junction length changes through time of cell B shown in a and 2i,j. Some junctions show correlation and change their length during the same time when others show anti-correlation. Each row and column represent a single junction and are color-coded as in a and 2I,J. Numbers represent the correlation coefficients of each pairs of junction compared. **(I)** Dot plot showing the correlation coefficients of junction length changes through time of junctions from multiple cells. Correlation of junctions shown in H are colored. **(J)** Plot showing the number of edges during the constriction period. Cells have different initial numbers of edges, and gradually lose them during the constriction period. **(K)** Plot showing the apical surface elongation index calculated by Tissue Analyzer, from 0 to 1, with 0 being perfectly round, and 1 corresponding to a line. Cells show fluctuation in their elongation during constriction, but exhibit no significant trend (trend showed by the linear regressions). **(L)** Plot showing apical surface orientation in the plane of the epiblast epithelial layer. Cells show fluctuation of orientation axis during constriction, and do not show significant pattern of elongation. 41 cells quantified from 4 embryos. *P<0.05 (Unpaired bilateral Mann-Whitney test). Error bars represent s.e.m. Scale bar, 10μm

**Figure 2-figure supplement 2:**
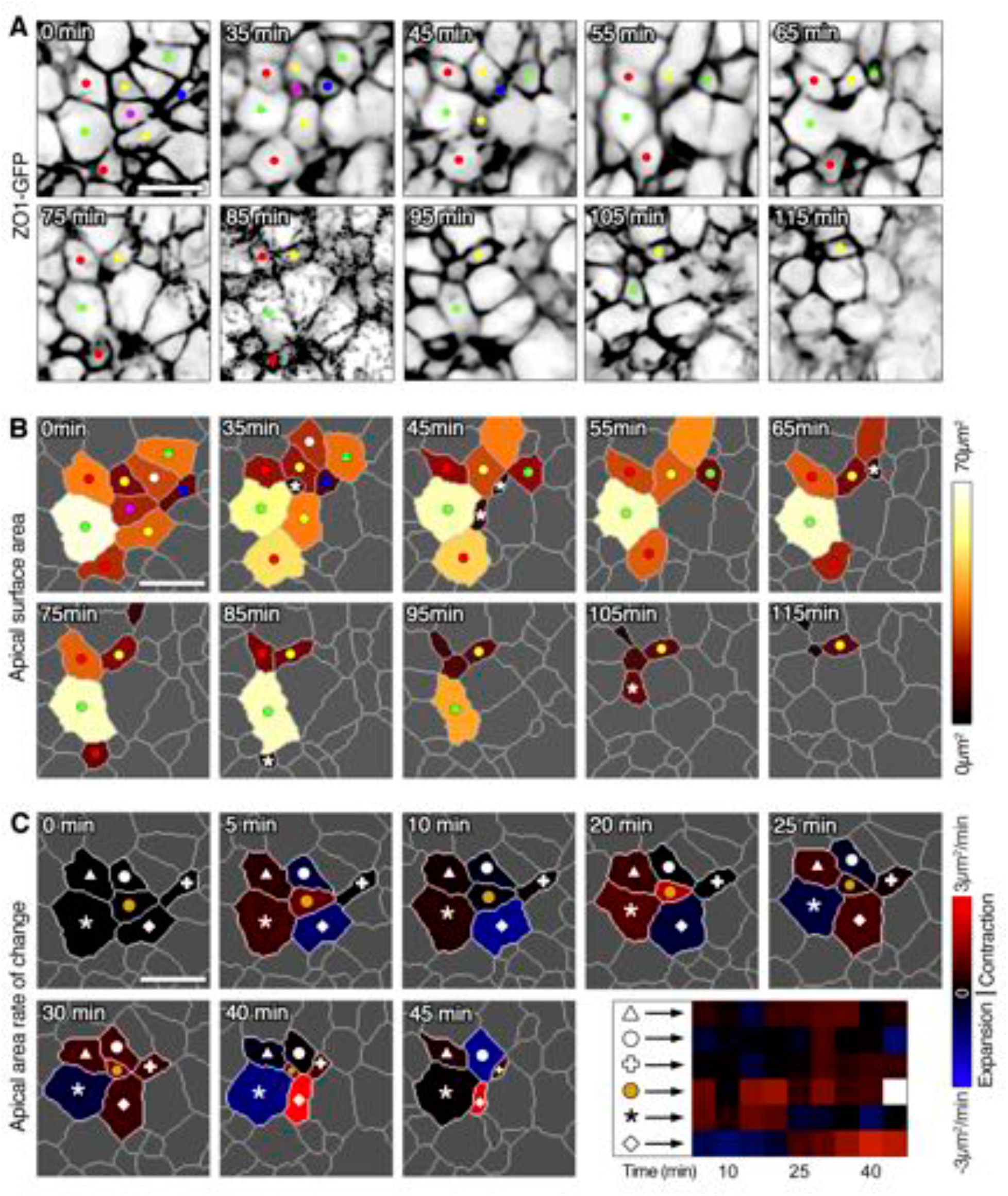
Asynchronous apical constriction at the multi-cellular level. **(A)** Time-series of ZO-1-GFP reporter expression identifying a cluster of epiblast cells exhibiting apparent stochastic apical constriction and cell ingression. **(B)** Membrane segmentation and color-coded time-series of apical surface area showing oscillation in apical surfaces in the cell cluster and asynchronous reduction of apical surfaces during ingression (asterisks). **(C)** Color-coded time-series and heat map plot of rate of change of apical surface area showing asynchronous oscillation in area and asynchronous constriction. Note that as one cell in the middle constricts and ingresses (orange dot), neighboring cells asynchronously change their apical surfaces; some will constrict and ingress shortly after, while others will only ingress at a later time point. Each row represents data for an individual cell. Scale bars, 10μm

**Figure 4-figure supplement 1:**
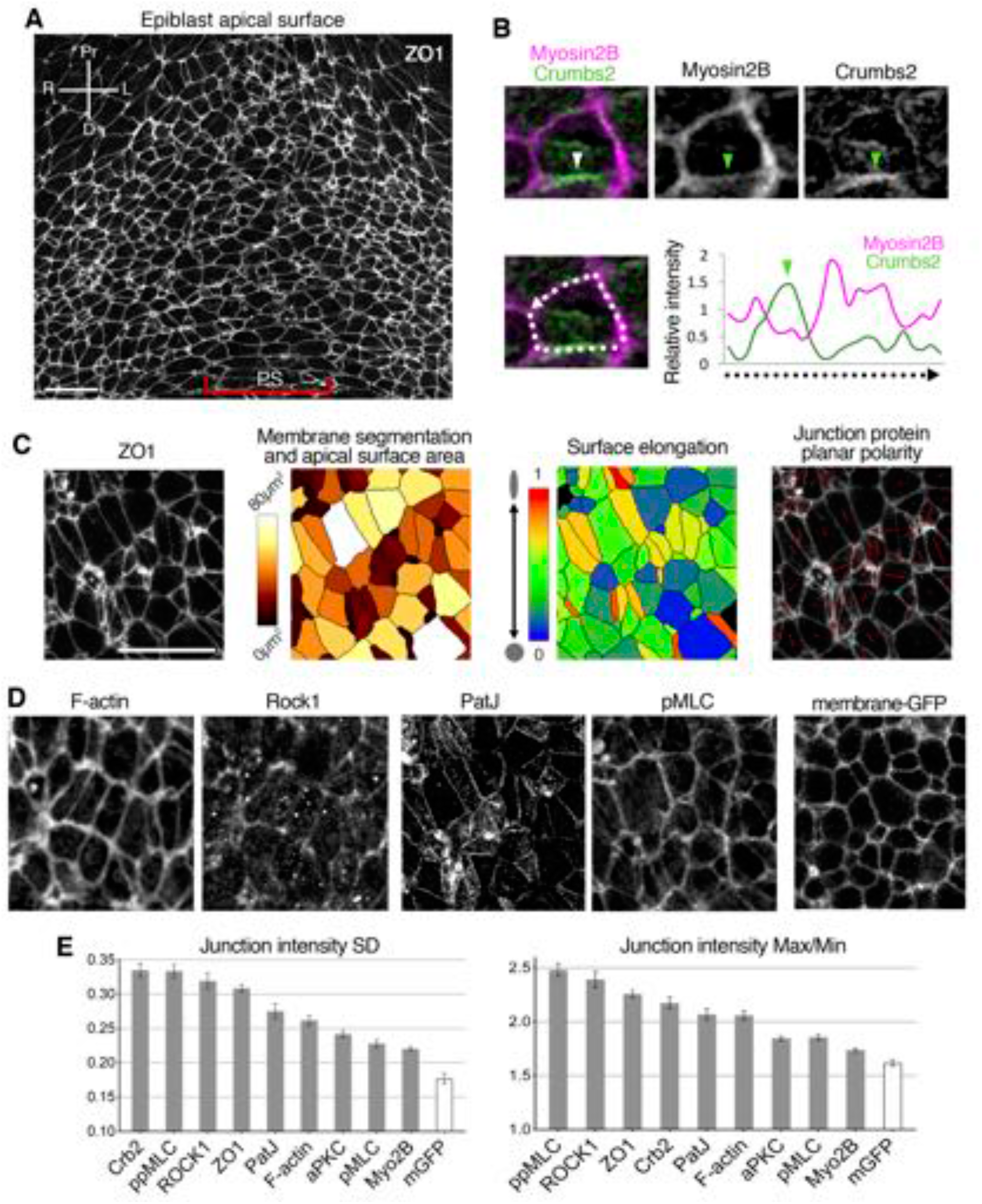
Complementary accumulation of Crumbs2 and Myosin2B, and anisotropy of apical components at the primitive streak. **(A)** ZO-1 immunostaining on fixed E7.5 embryo showing the apical surface of epiblast cells in the posterior. **(B)** High magnification view of a single cell stained for Myosin2B (magenta) and Crumbs2 (green) showing strong Crumbs2 accumulation on one junction (arrowhead) and predominant Myosin accumulation on other junctions. White dashed line and arrow indicate the membrane intensity measurement shown in the plot. The junction with the highest level of Crumbs2 (arrowhead) has the lowest level of Myosin, whereas Myosin predominates (and Crumbs is reduced) at the rest of the junctions. **(C)** ZO-1 immunostaining used for cell apical membrane identification and segmentation, allows quantification of junctional intensity of staining, and determination of a number of parameters, including apical surface area, surface elongation and protein planar polarity. **(D)** Staining F-actin, Rock1, PatJ and pMLC exhibiting different patterns of accumulation at apical cell-cell junctions, and a membrane-GFP reporter more homogeneously present at the apical membrane. **(E)** Plots of anisotropy parameters, junctional Standard Deviation and Max/Min. Proteins exhibit a range of distributions and anisotropy levels, some like Crumbs2 and ppMLC show a high level of anisotropy, others like PatJ and F-Actin show moderate anisotropy, and some like aPKC pMLC and Myosin2B show a lower level of anisotropy closer to the level of the negative control membrane-GFP. Error bars represent s.e.m.

**Figure 4-figure supplement 2:**
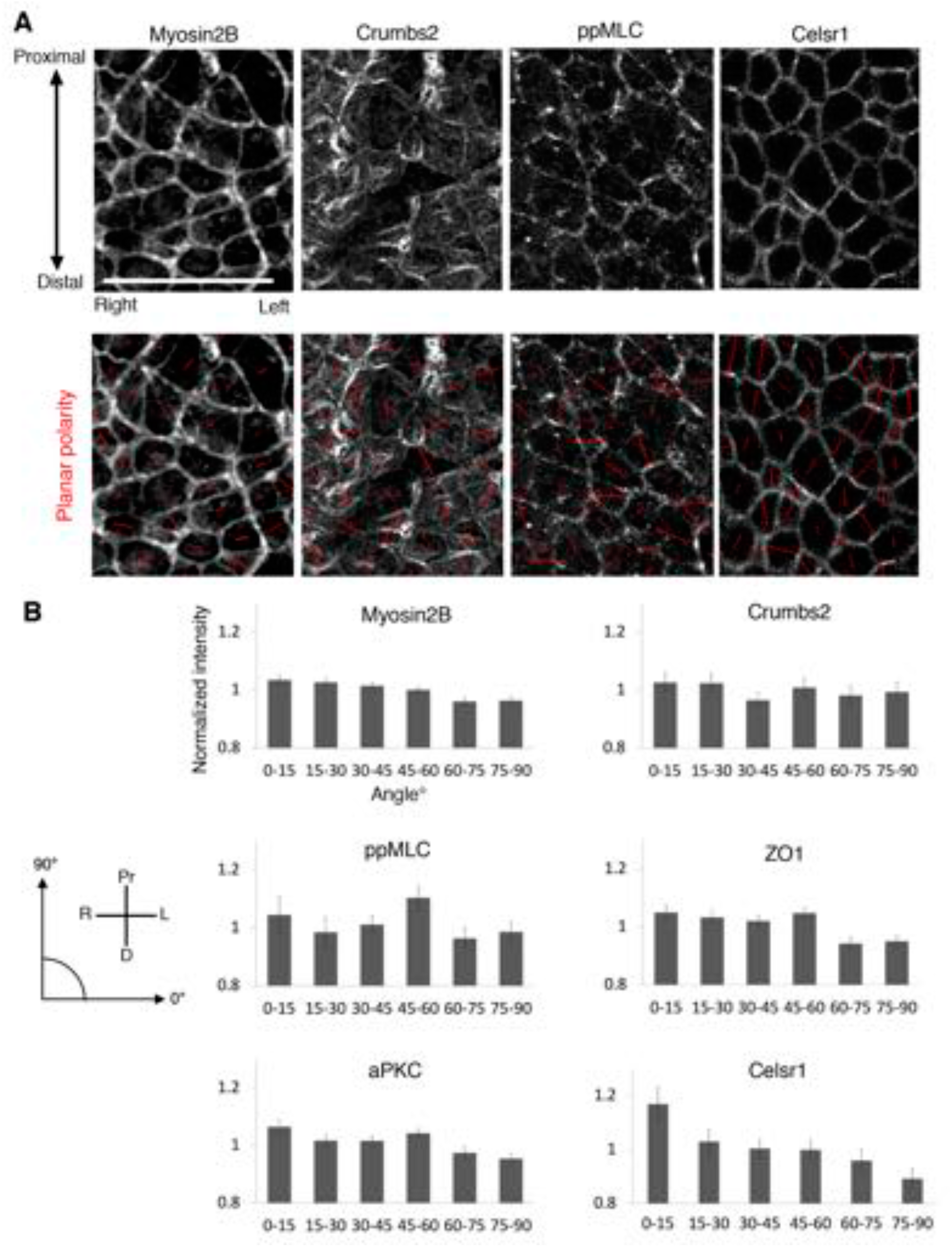
Apical proteins show anisotropy accumulation but no planar polarity. **(A)** Immunostaining for Myosin2B, Crumbs2, ppMLC and Celsr1 with associated polarity visualization (red tensor) calculated after membrane segmentation, showing the orientation (orientation of the tensor) and amplitude of polarity (length of the tensor). **(B)** Junctional protein intensity plotted as a function of junctional angle. 0° is parallel to the left/right axis and 90° is parallel to the proximal/distal axis. The apical proteins described previously and measured here (Myosin2B, Crumbs2, ppMLC, ZO-1 and aPKC) do not show planar polarity, red bars in A showing polarity of intensity are not oriented in specific axis, and intensity in B show no bias at a particular angle. By contrast, the PCP protein Celsr1 exhibits evident polarity, red bars tend to be vertically oriented, and intensity in B shows a significant increase in Celsr1 levels at edges oriented parallel to the left/right axis. Celsr1 preferentially accumulates at horizontal junctions (left-right embryo orientation). Error bars represent s.e.m.

**Figure 5-figure supplement 1:**
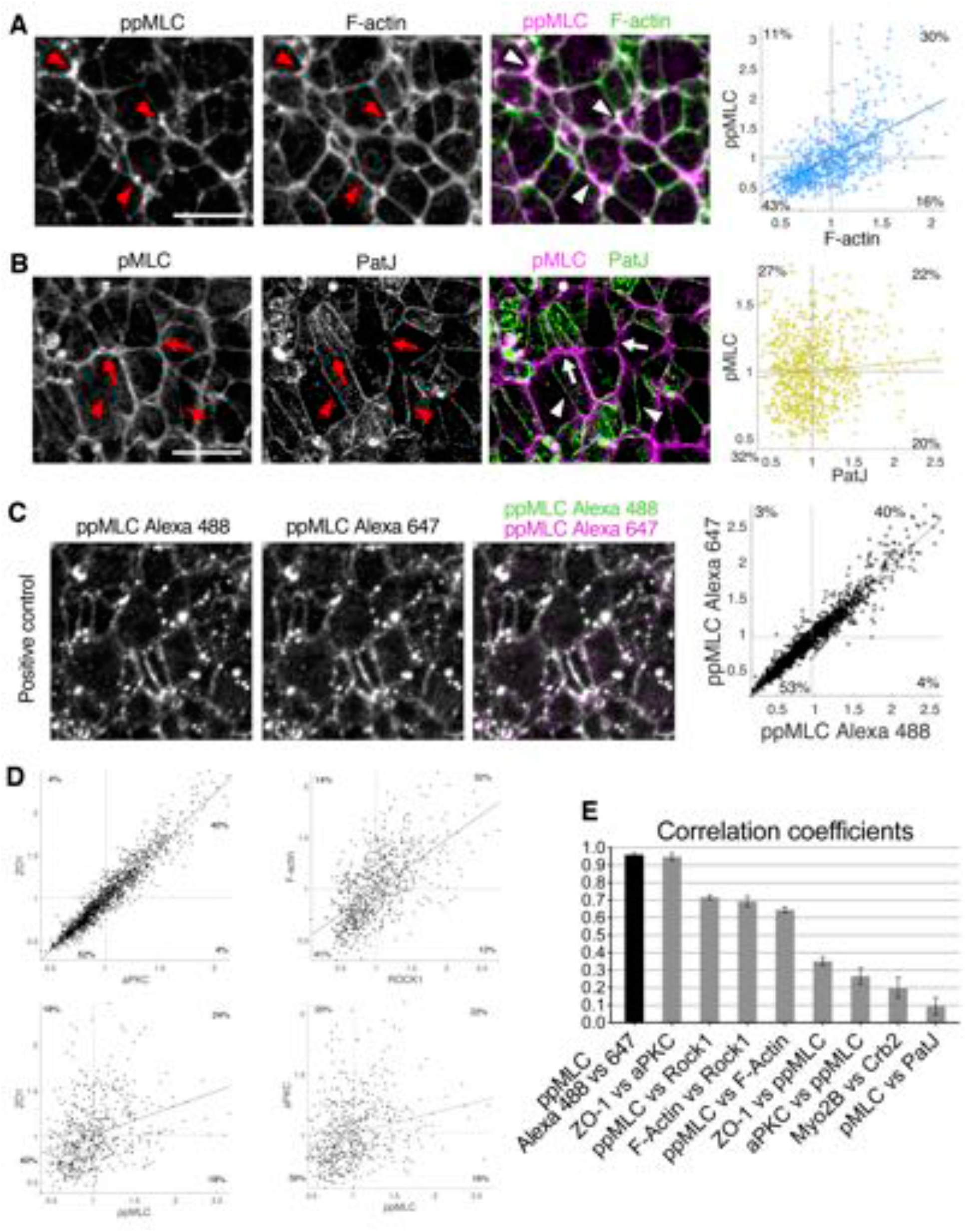
Correlation of protein distributions at apical junctions. **(A)** Co-immunostaining of ppMLC and F-Actin reveals a tendency to accumulate on the same junctions. Note colocalization of fluorescent signal on most junctions (arrowheads). Scatter plot on the right showing elevated intensity correlation of the two proteins. Each point represented the mean intensity at a single junction. **(B)** Co-immunostaining of pMLC and PatJ reveals that they tend to accumulate on different junctions. Note junctions where Myosin accumulation predominates (arrows), and junctions where Crumbs2 predominates (arrowheads). Corresponding scatter plot shows more spread distribution and no correlation. **(C)** Immunostaining for ppMLC simultaneously revealed by two distinct secondary antibodies (Alexa 488 and 647) as positive control, and corresponding scatter plot showing strong junctional intensity correlation after membrane segmentation. **(D)** Scatter plots showing the correlation of intensity of other pairs of proteins. **(E)** Correlation coefficients associated with scatter plots shown in A,B,C,D and Figure 5. Error bars represent s.e.m.

**Figure 5-figure supplement 2:**
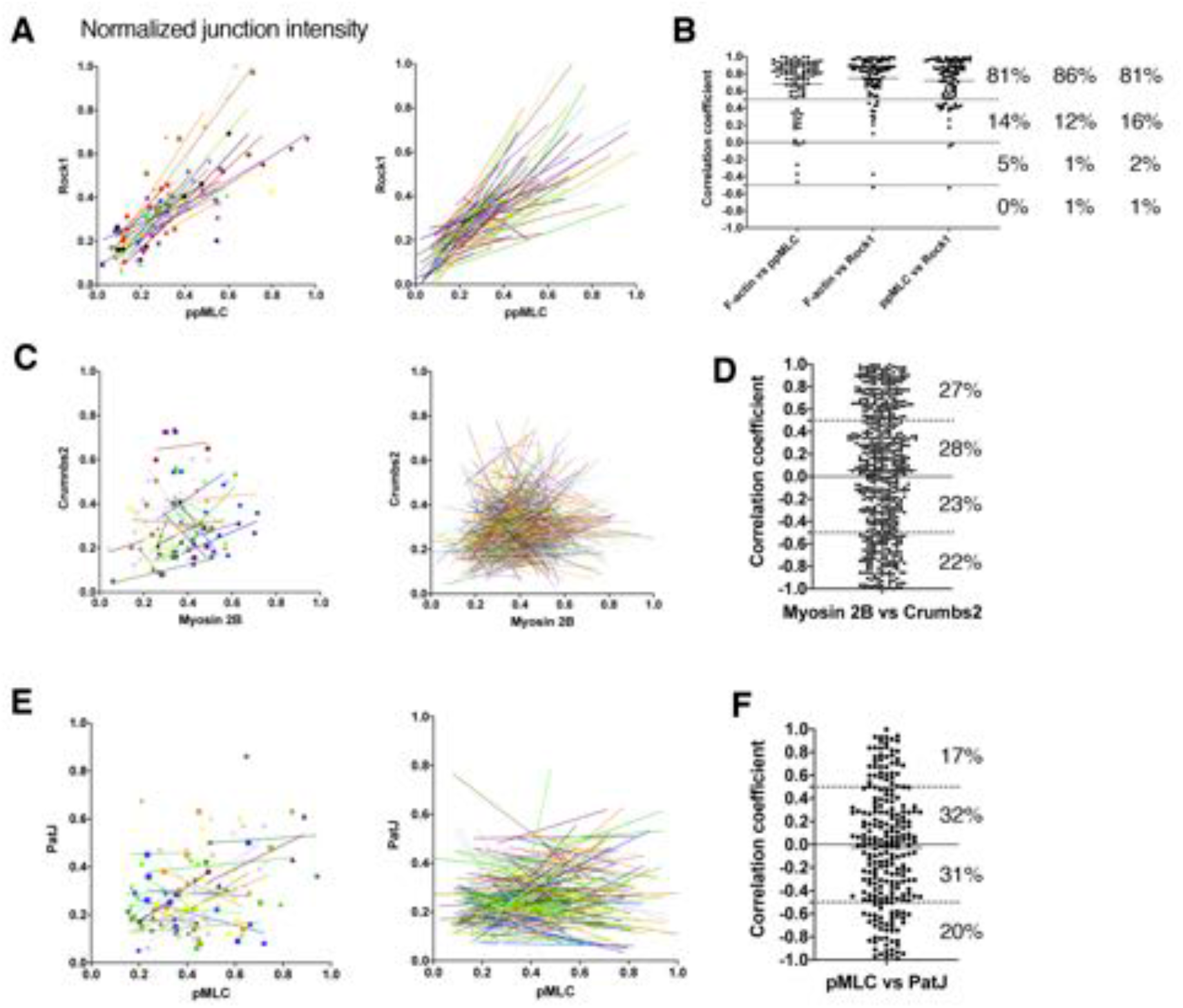
Junctional correlation of proteins at the single cell level. **(A)** Example of scatter plot showing the junctional intensity correlation of ppMLC and Rock1 in a cohort of cells. Each symbol represents one junction, grouped by cells (color coded) with respective correlation trends. Plot on the right shows only the correlation trend for a larger number of cells. **(B)** Correlation coefficients of ppMLC and Rock1 from plot shown in A, as well as F-actin and ppMLC, and F-actin and Rock1. Each dot represents the coefficient of correlation of junctions intensity of a single cell. % show the distribution of correlation coefficients in the different quarters. **(C)** Example of junctional intensity correlation of single cells for Crumbs2 and Myosin2B, depicting a mix of cells with correlation, some with no correlation, and some with anti-correlation. **(D)** Correlation coefficients reflecting the scatter plot shown in C, showing a broad distribution, from correlation (+1) to anti-correlation (−1). % show the distribution of correlation coefficients. **(E)** Example of junctional intensity correlation of single cells for PatJ and pMLC, depicting a mix of cells with correlation, some with no correlation, and some with anti-correlation. **(F)** Correlation coefficients reflecting the scatter plot shown in C, showing a broad distribution, from correlation (+1) to anti-correlation (−1). % show the distribution of correlation coefficients.

**Figure 6,7-figure supplement 1:**
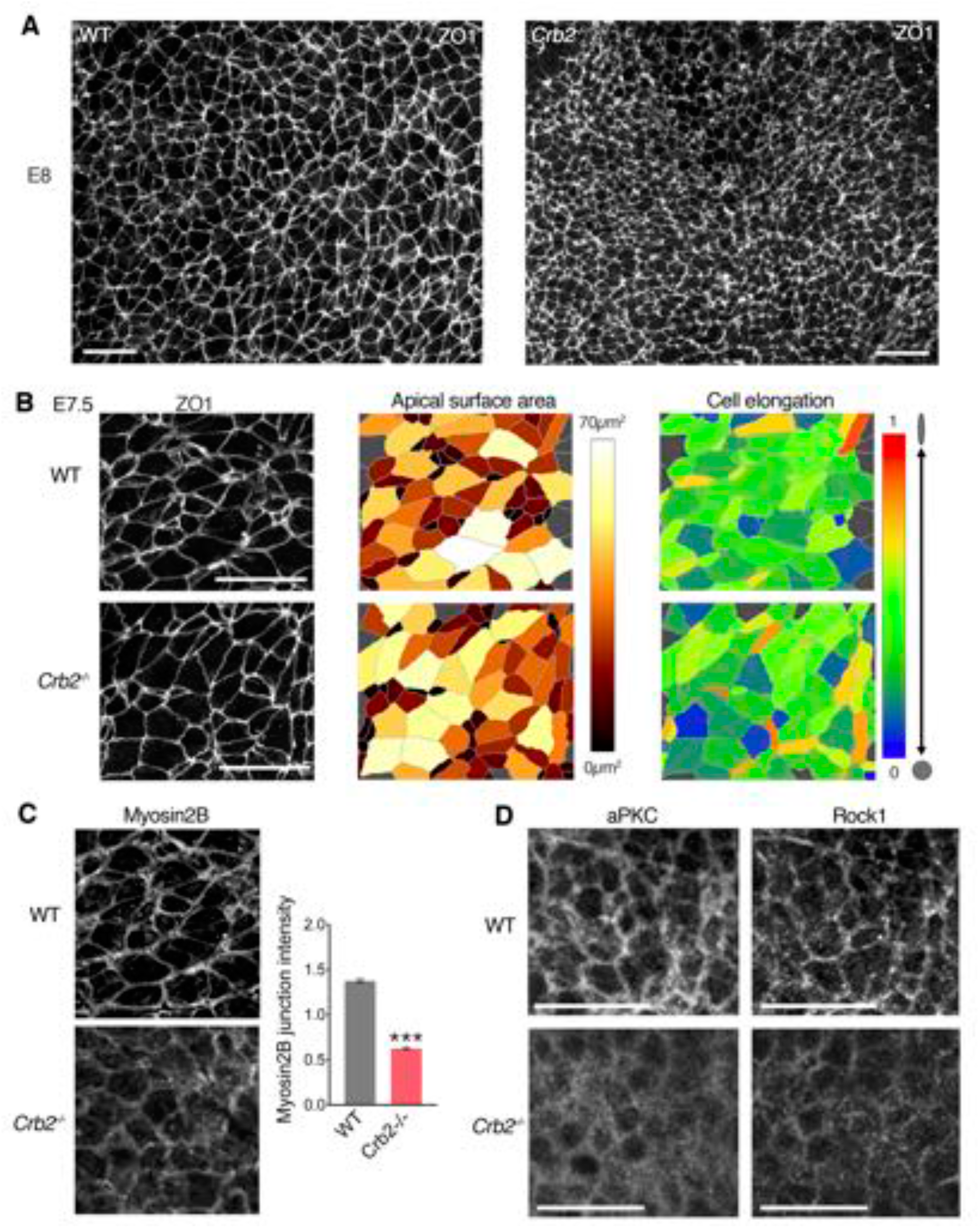
Cellular and molecular defects in *Crumbs2* mutant embryos. **(A)** Low magnification of ZO-1 staining show in the posterior epiblast in WT and *Crumbs2* mutants at E8.0. **(B)** ZO-1 immunostaining showing the apical surface of WT and *Crumbs2* mutant embryos at E7.5. Segmentation and color code of apical surface areas and surface elongation reveal normal size and shape of apical surface area in *Crumbs2* mutants at E7.5. **(C)** Immunostaining for Myosin2B at E7.5, showing reduced intensity in *Crumbs2* mutant embryos compared to WT. Graph shows significantly reduced Myosin2B intensity at junctions. **(D)** Immunostaining for aPKC and Rock1 in WT and *Crumbs2* mutants at E7.5 (intensity quantifications shown in Figure 7C). ***P<0.001 (Unpaired bilateral Mann-Whitney test). Error bars represent s.e.m.

**Video S1**. Time-lapse imaging of a membrane-GFP reporter (recombined *mT/mG*) expressing embryo at embryonic day E7.5, showing apical constriction (right panel) associated to ingression, as cells move out of the epiblast epithelial layer basally to join the underlying mesoderm (left panel). Images were denoised and deconvolved, and represent single planes of optical sections and apical cell surfaces. The time-lapse spans 70min with 10min interval between frames.

**Video S2**. Time-lapse imaging of the posterior of a ZO-1-GFP reporter expressing embryo in the vicinity of the primitive streak at embryonic day E7.5. Tracking of individual cells reveals lateral epiblast cells converging toward the midline prior to their ingression. Cells around the midline (middle row) do not exhibit extensive movement in the plane of the epithelium. Movie of 240min with 5min interval between frames.

**Video S3**. Time-lapse of ZO-1-GFP showing rearrangement of cells in the epiblast near the primitive streak (top panel, 125min), and cell apical constriction and ingression away from the primitive streak, as the epiblast flows (to the left side) toward the midline (bottom panel, 55min). 5min interval between frames.

**Video S4**. Time-lapse of ZO-1-GFP highlighting 3 cells apically constricting and ingressing in the primitive streak region. 60min total with 5min interval between frames.

**Video S5**. Time-lapse of ZO-1-GFP highlighting a single cell at the primitive streak apically constricting. Membrane segmentation is performed over time such that different parameters can be followed and quantified, including the apical surface area, a cell’s elongation and orientation, and the number of junctions. 95min total with 5min interval between frames.

**Video S6**. Segmentation of time-lapse showing color-code of cell apical surface (left panel), constriction rate (middle panel) and junction length (right panel) of a cell A shown in Figure 2. 40min total with 5min interval between frames.

**Video S7**. Time-lapse of ZO-1-GFP showing a cell cluster at the primitive streak. Most of the cells gradually constrict and ingress, and cells tracked after segmentation and color-coded for apical surface area and changes in surface area. Images were denoised and deconvolved. 115min with 5min interval between frames.

